# Beneficial reversal of dominance maintains a resistance polymorphism under fluctuating insecticide selection

**DOI:** 10.1101/2024.10.23.619953

**Authors:** Marianthi Karageorgi, Anastasia S. Lyulina, Mark C. Bitter, Egor Lappo, Sharon I. Greenblum, Zach K. Mouza, Caitlynn T. Tran, Andy V. Huynh, Hayes Oken, Paul Schmidt, Dmitri A. Petrov

**Author notes:** Corresponding authors: M.K., P.S., and D.A.P. These authors contributed equally (Paul Schmidt, Dmitri A. Petrov).

## Abstract

Large-effect functional genetic variation is commonly found in natural populations, even though natural selection should erode such variants. Theory suggests that under fluctuating selective pressures, beneficial reversal of dominance - where alleles are dominant when beneficial and recessive when deleterious - can protect these loci from selection, allowing them to persist. However, empirical evidence for this mechanism remains elusive because testing requires direct measurements of selection and dominance in natural conditions. Here, we show that insecticide-resistant alleles at the *Ace* locus in *Drosophila melanogaster* persist worldwide at intermediate frequencies and exhibit beneficial reversal of dominance. By combining laboratory and large-scale field mesocosm experiments with insecticide manipulation, and mathematical modeling, we show that the benefits of the resistant *Ace* alleles are dominant while their fitness costs recessive. We further show that fluctuating insecticide selection generates chromosome-scale genomic perturbations at sites linked to the resistant *Ace* alleles, revealing broader genomic consequences of this mechanism. Overall, our results suggest that beneficial reversal of dominance contributes to the maintenance of functional genetic variation and impacts patterns of genomic diversity via linked fluctuating selection.

## Main text

Large-effect functional variants responding to temporally fluctuating selective pressures persist in natural populations across taxa, from ancient to recently evolved polymorphisms to anthropogenic pressures^1,2^. A striking example comes from studies of seasonal dynamics in *Drosophila*, where alleles can change frequencies by tens of percent per season^3–6^ yet remain polymorphic over millions of generations^3^. However, their maintenance is puzzling because population genetics theory suggests that the conditions for preserving functional genetic variation under temporally fluctuating selection are restrictive^7^. One mechanism that can relax these conditions is beneficial reversal of dominance^8,9^, where alleles are partially or completely dominant when beneficial in favorable environments but become partially or completely recessive when deleterious in unfavorable environments^10,11^. This mechanism acts as a stabilizing force, protecting recessive deleterious alleles from selection while enabling rapid adaptation when selection shifts. An extension of Wright’s physiological theory of dominance^12^ further suggests that this reversal can occur when alternative adaptive alleles at a locus function as loss-of-function variants in their unfavored environments^10^. However, empirical evidence for the mechanism remains scarce because testing requires direct measurements of selection and dominance of adaptive alleles on fitness in natural conditions.

The *Ace* locus in insects provides an ideal genetic system to test whether beneficial reversal of dominance can maintain functional variation under fluctuating selection^13^. *Ace* encodes acetylcholinesterase (AChE), a key enzyme in the insect nervous system responsible for breaking down acetylcholine^13,14^. Organophosphate insecticides, introduced in the 1950s and 1960s, act as irreversible inhibitors of AChE, causing insect death by binding to its active site and preventing removal of acetylcholine from the synapses upon firing^15^. These insecticides have imposed seasonally fluctuating selection pressures on resistant insect populations, as seen in seasonal resistance fluctuations in field studies^16–18^. Following Wright’s physiological theory of dominance, resistant alleles should be loss-of-function and thus recessive in pesticide-free environments, while sensitive alleles should become loss-of-function and thus recessive in pesticide-rich environments^19,20^. Consistent with this prediction, previous studies across resistant insect populations suggest that resistant *Ace* alleles follow patterns of beneficial reversal of dominance often showing dominant benefits and/or recessive costs for fitness-associated traits^19,21^, though direct evidence is lacking.

Here, we test whether beneficial reversal of dominance under temporally fluctuating selection contributes to the maintenance of the resistant *Ace* alleles in natural populations of *D. melanogaster*, and investigate the broader genomic consequences of the mechanism. In *Drosophila melanogaster*, resistant *Ace* alleles have repeatedly evolved in natural populations in response to organophosphate selection^22,23^, and enzymatic assays suggest the resistance mutations are beneficial under insecticide exposure and costly in its absence^22,24^. However, the phenotypic and fitness consequences of these mutations have not been studied. We combine population genomic analysis, laboratory assays, and a novel field mesocosm system that enables real-time tracking of genome-wide allele frequencies and resistance under controlled insecticide manipulation^5^ to test for beneficial reversal of dominance of the resistant *Ace* alleles. The field mesocosm system establishes a direct experimental approach to observe beneficial reversal of dominance and its genomic consequences in response to controlled selective pressures in semi-natural conditions.

### Resistant *Ace* alleles persist in natural populations and respond to fluctuating insecticide selection

We first examined whether resistant *Ace* alleles exhibit signatures consistent with beneficial reversal of dominance under temporally fluctuating selection in natural *D. melanogaster* populations. This mechanism predicts that resistant *Ace* alleles should fluctuate in response to temporally fluctuating selection while persisting at intermediate frequencies. Previous studies identified resistant *Ace* alleles in *D. melanogaster* with mutations at four conserved sites near the AChE active site (I161V, G265A, F330Y, G368A), with <u>V</u>GFG and <u>VAY</u>G as the most common resistance haplotypes^22,23^ (inset, **Fig. 1a**). Our analysis of inbred lines from Pennsylvania (LNPA^5^) and North Carolina (DGRP^25^) confirmed these as the most common resistant haplotypes in resistant *Ace* alleles (**Extended Data Fig. 1a, b**), with the LNPA also carrying <u>V</u>GF<u>A</u> at ∼13% frequency. Because I161V is present across the most common haplotypes, we used it as a shared marker to track resistant *Ace* alleles in natural populations.

**Figure 1.**
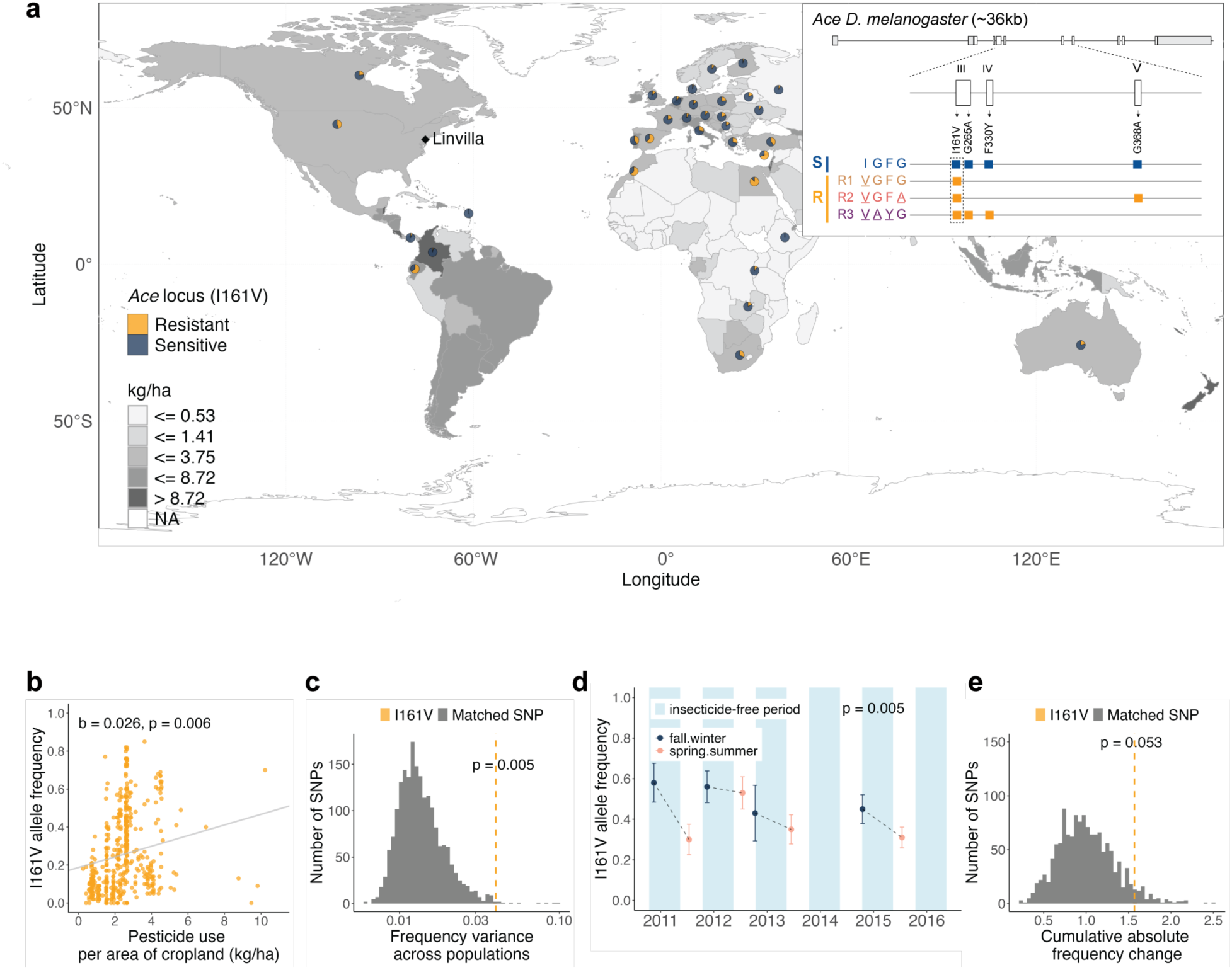
Organophosphate-resistant *Ace* alleles persist at intermediate frequencies in natural *Drosophila melanogaster* populations and respond to spatially and temporally fluctuating insecticide selection. **a**, Allele frequency map of I161V (pie charts), a marker of organophosphate-resistant *Ace* alleles, overlaid on insecticide use intensity (shading). Inset shows common resistant *Ace* alleles with three different resistance haplotypes (R1: <u>V</u>GFG, R2: <u>V</u>GF<u>A</u>, R3: <u>VAY</u>G) in *D. melanogaster* (See also **Extended Data Fig. 1** for frequencies of resistant haplotypes in DGRP and LNPA panels of fully sequenced inbred lines). **b,** I161V frequency increases with increased pesticide use (linear mixed-effect model). **c,** I161V shows higher variance in frequency compared to matched control SNPs across populations (orange dashed line indicates I161V). **d,** Seasonal changes in I161V frequency at Linvilla Orchards showing decline from fall (dark blue) to spring (pink) samples during insecticide-free periods (turquoise, mid-fall to mid-spring) (generalized linear model, mean ± s.e.m.). **e,** Cumulative frequency changes at Linvilla Orchards samples show elevated allele frequency fluctuations of I161V compared to matched control SNPs (orange dashed line indicates I161V).

While resistant *Ace* alleles experience temporally fluctuating selection, they are also subject to spatial variation in insecticide exposure. To disentangle these effects, we analyzed genomic data from worldwide *D. melanogaster* populations from the Drosophila Evolution Over Space and Time (DEST) database^26^ and country-level pesticide use data from the Food and Agricultural Organization of the United Nations (FAO)^27^ (see **Methods f**or details). Spatially, I161V frequency increases with pesticide use intensity across countries (linear mixed-effect model, p = 0.006) (**Fig. 1a, b**) and shows stronger spatial variation compared to control SNPs matched for recombination and average frequency (p = 0.005) (**Fig. 1c**). Temporally, seasonal sampling at Linvilla Orchards, a non-organic farm where samples were collected in spring and fall from 2009 to 2015, revealed a consistent fall-to-spring decline in I161V frequency (generalized linear model, p = 0.005), suggesting either fitness costs of the resistant *Ace* alleles in winter (insecticide-free time period) or seasonal migration (**Fig. 1d**). In addition, the I161V polymorphism showed stronger temporal fluctuations than matched control SNPs (p = 0.053) (**Fig. 1e**). Therefore, these findings are consistent with beneficial reversal of dominance, showing that resistant *Ace* alleles persist at intermediate frequencies under temporally fluctuating selection.

### Lab experiments reveal dominant benefits and recessive (or co-dominant) costs of resistant *Ace* alleles for fitness-associated phenotypes

We then examined whether the resistant *Ace* alleles exhibit beneficial reversal of dominance for fitness associated phenotypes relevant to organophosphate-rich and -free environments. In organophosphate-rich environments, resistance is primarily under selection and the resistant *Ace* alleles should be beneficial because the resistance mutations alter the shape of AChE’s active site to prevent pesticide binding^22^. In organophosphate-free environments, life history traits are primarily under selection and the resistant *Ace* alleles should be deleterious because the resistance mutations impair AChE activity and stability^24^. Following Wright’s theory of dominance, the resistant alleles should be dominant in organophosphate-rich environments where sensitive alleles become loss-of-function, and recessive in organophosphate-free environments where they themselves become loss-of-function^20^ (**Fig. 2a,b**). The underlying basis of this mechanism is that even one copy of the functional allele in each favored environment should provide sufficient gene activity to carry out the required function.

**Figure 2.**
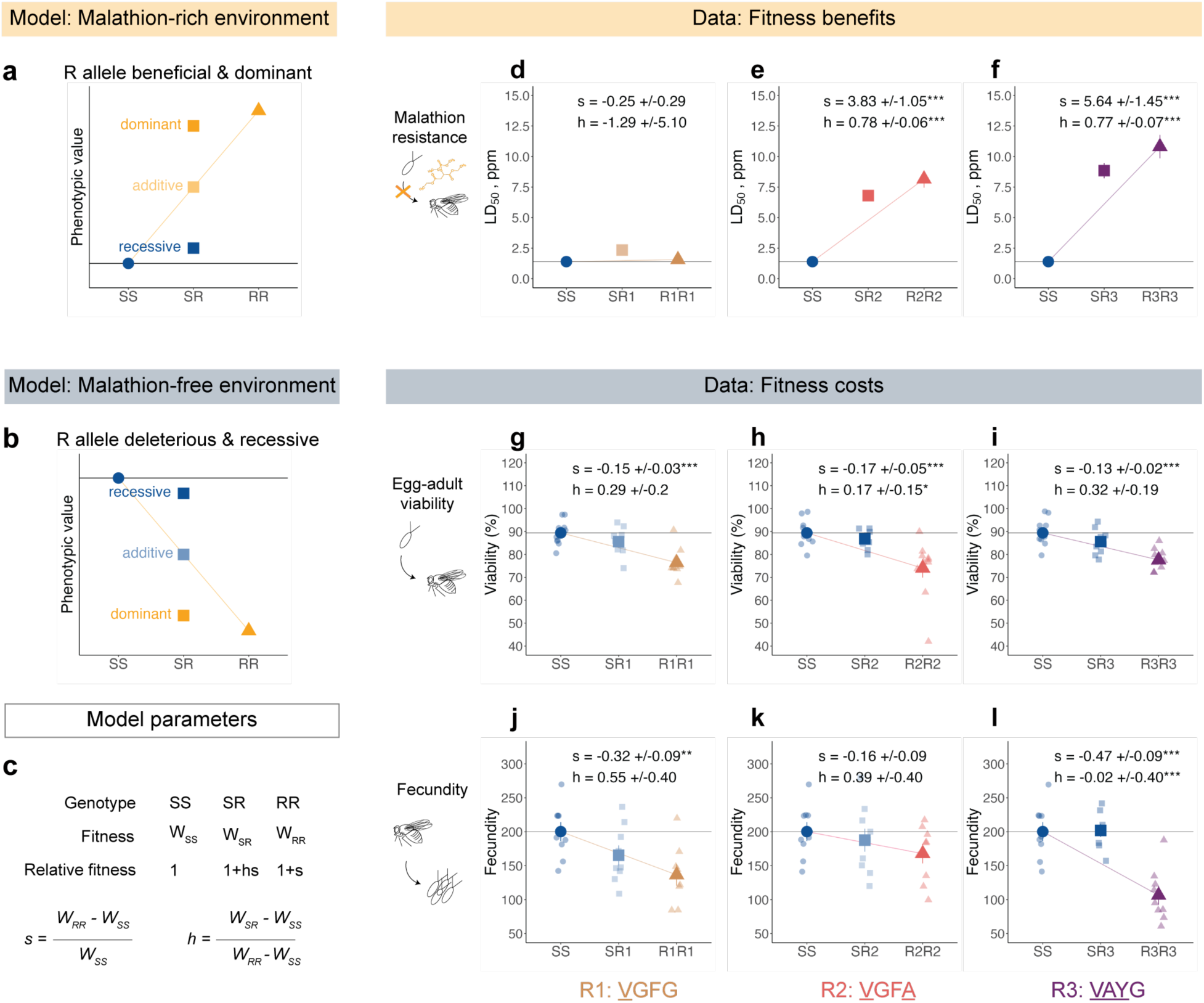
Beneficial reversal of dominance of organophosphate-resistant *Ace* alleles for fitness-associated phenotypes relevant to malathion-rich and malathion-free environments. **a-b**, Under the beneficial dominance reversal, R alleles should be beneficial and partially or completely dominant in malathion-rich environments (*W_SR_* > (*W_SS_*+*W_RR_*)/2, orange), and deleterious and partially or completely recessive in malathion-free environments (*W_SR_* < (*W_SS_*+*W_RR_*)/2, blue). **c,** Selection (*s*) and dominance (*h*) coefficients for each fitness-associated phenotype are estimated based on the viability selection model. If s>1, the resistant R *Ace* allele is beneficial; if s<1, it is deleterious. If h=1, the resistant R *Ace* allele is dominant to the sensitive S allele; if h=0, the reverse is true. **d-l**, Comparison of malathion LD_50_ (**d,e,f**), egg-adult viability (n=10 technical replicates, 50 eggs/replicate) (**g,h,i**), and fecundity (n=8–10 technical replicates, 5 females/bottle/3 days) (**j,k,l**) among *Ace* genotypes. Malathion LD_50_ was calculated from a four-parameter logistic model fitted to egg-adult viability measurements on media with a range of malathion concentrations (n=2 technical replicates, 50 eggs/replicate, see **Extended Data Fig. 2**). Egg-adult viability and fecundity data were analyzed with logistic and linear regression models.

To test these predictions, we used the LNPΑ panel of inbred lines (**Extended Data Fig. 1a,b**) to generate recombinant populations with homozygous sensitive, homozygous resistant, and heterozygous genotypes for each resistant allele (S: IGFG, R1: <u>V</u>GFG, R2: <u>V</u>GF<u>A</u>, R3: <u>VAY</u>G), an approach^28^ that controls for the genomic background, and then measured resistance to the organophosphate insecticide malathion and two key life history traits, viability (egg to adult survival) and fecundity (number of eggs laid by gravid females) (see **Methods** for details). For malathion resistance, the R2 and R3 alleles significantly increased LD_50_ (dose causing 50% egg-to-adult mortality) by ∼5.8x and ∼7.8x respectively (p < 0.001), and were strongly dominant over the sensitive S allele, with dominance *h* of 0.78 and 0.77 for R2 and R3, respectively (p < 0.001 for h > 0.5 in both cases). The R1 allele showed no significant resistance (p = 0.646) with inconclusive dominance effects (p = 0.751) (**Fig. 2, Extended Data Fig. 2, Extended Data Table 1**). For viability and fecundity, all homozygous resistant genotypes exhibited lower phenotypic fitness values compared to the homozygous sensitive genotype, for either or both of them, with viability reductions ranging from 12.4% to 16.9% (all p < 0.001) and fecundity reductions from 16.2% (R2R2, p = 0.37) to 46.8% (R3R3, p < 0.001). The costs driven by the three resistant alleles were either significantly recessive for at least one of the phenotypes for the strongest resistance alleles R2 and R3 (viability R2: h = 0.17, p = 0.027, fecundity R3: h = -0.02, p = 0.001), or indistinguishable from codominance for R1 (**Fig. 2g-l, Extended Data Tables 2-3**). Therefore, these findings support the beneficial reversal of dominance of the resistant *Ace* alleles for key fitness associated phenotypes in organophosphate-rich and -free environments.

### Field mesocosm experiments reveal dominant benefits and recessive costs of resistant *Ace* alleles for fitness

We next tested whether beneficial reversal of dominance of the resistant *Ace* alleles drives evolutionary responses to fluctuating insecticide selection in semi-natural conditions and examined its genomic consequences. Given the dominant benefits and recessive costs observed in the laboratory experiments, we predicted a rapid increase in the frequency of the resistant *Ace* alleles and in the level of overall resistance at insecticide concentrations similar to the LD_50_ of the sensitive homozygous genotype. We also predicted a frequency-dependent decline of the resistant *Ace* alleles in the absence of insecticide because recessive costs become exposed in resistant homozygotes when the resistant *Ace* alleles are at high frequencies. For the genomic consequences, we expected that large-amplitude fluctuations of the resistant *Ace* alleles due to fluctuating insecticide selection would drive significant effects on linked genomic variation, as theoretical models suggest^29^. These effects on linked sites should extend across large genomic distances, with their magnitude scaling with the rate of adaptive allele frequency shifts^30^.

To test these predictions, we conducted a field mesocosm experiment to track the evolutionary dynamics of the resistant *Ace* alleles under controlled seasonal insecticide selection (**Fig. 3a**, see **Methods**). Using an experimental orchard in Pennsylvania, we established 20 replicate *D. melanogaster* populations derived from a baseline population constructed from the LNPA panel of inbred lines on June 21, 2021 (**Extended Data Fig. 1**). These lines were outbred for 4 generations in large indoor cages before ∼1000 males and females were used to seed each outdoor cage (see **Methods**). After ∼4 weeks of untreated propagation, we divided the populations into ten malathion-treated and ten untreated outdoor cages. To mimic natural insecticide exposure, we applied stepwise increasing concentrations of commercial malathion in the growth media of the treated populations (1.5–2.5 ppm from July 21 to August 24, followed by 7.5 ppm until September 6, 2021), exerting strong selection for resistant *Ace* alleles based on our lab LD_50_ measurements (**Extended Data Fig. 2**). Following insecticide removal, all populations were maintained without malathion through December 22, 2021. Throughout the experiment, we tracked population dynamics, malathion resistance by estimating LD_50_, as well as trajectories of resistant *Ace* alleles and linked sites in response to fluctuating malathion treatment at eight timepoints (∼every one to two generations) (see **Methods**).

**Figure 3.**
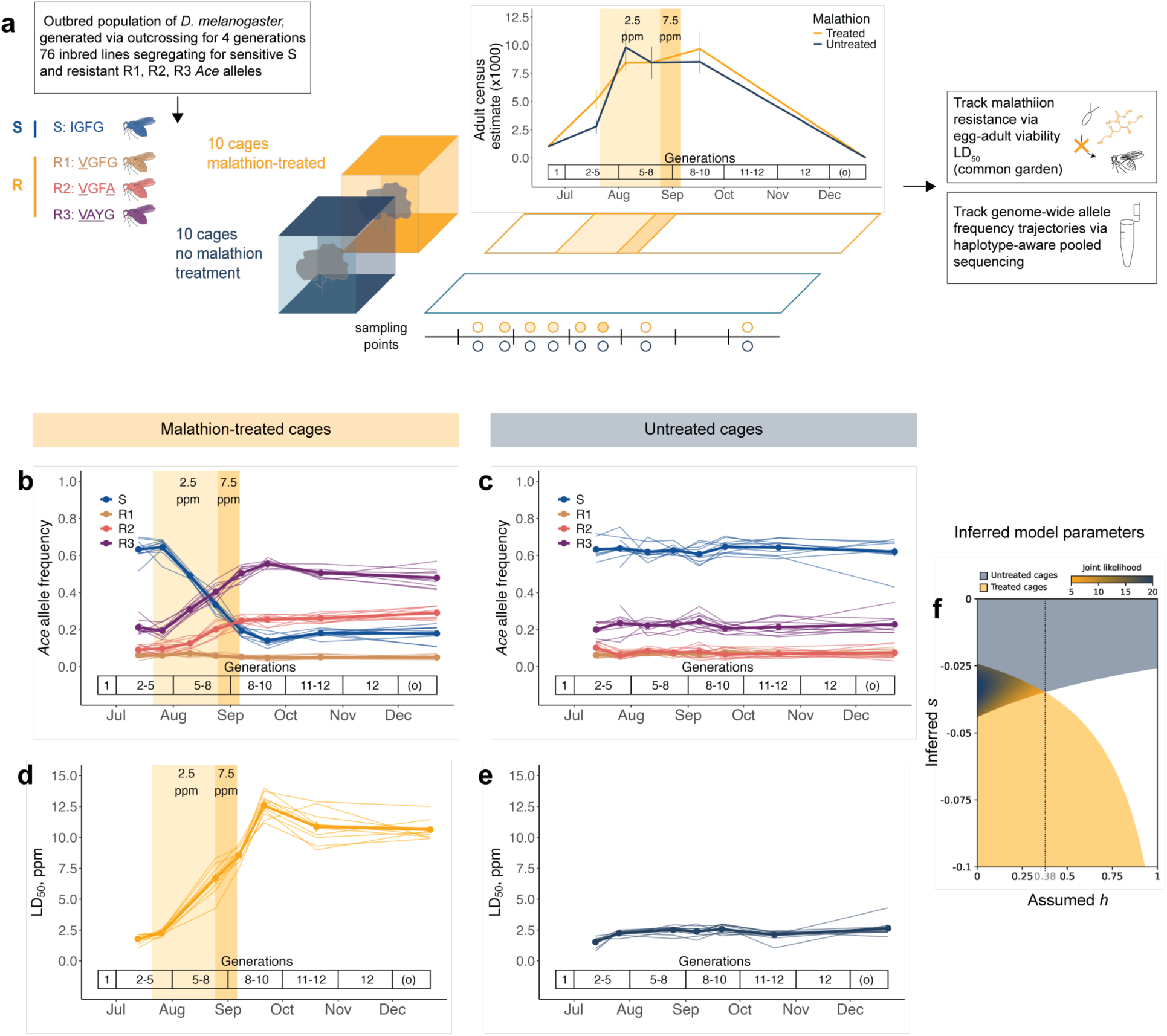
Beneficial reversal of dominance of resistant *Ace* alleles for fitness in an ecologically realistic environment. **a,** Twenty field mesocosm cages were seeded with an outbred population of *D. melanogaster* generated via outcrossing for 4 generations 76 inbred lines carrying the sensitive S and resistant R1, R2, and R3 alleles. Ten cages received a malathion pulse after the initial population establishment. Eight timepoints were sampled to track the evolution of malathion resistance and genome-wide allele in the treated and untreated populations. Malathion did not affect estimated adult fly population sizes (mean ± s.e.m.) in treated populations. **b-c,** Trajectories of *Ace* alleles in treated (**b**) and untreated (**c**) populations. In treated populations, the malathion pulse led to a significant increase in the frequency of the resistant R2 and R3 alleles, followed by a decline in the frequency of R3. *Ace* allele frequencies remained stable in the untreated populations. Light lines/points show individual allele trajectories, dark lines/points represent the mean ± s.e.m. (n = 10 populations per treatment group). **d-e,** The malathion pulse led to a significant increase and subsequent decline of resistance (LD_50_, ppm) in the treated populations while resistance remained stable in the untreated populations. For LD_50_ calculations see **Methods** and Light lines/points show individual populations (orange, treated; blue, untreated), dark lines/points represent mean ± s.e.m. (n=10 populations per treatment group).**f,** Inferred selection (s) and dominance (h) coefficients from changes in the resistant *Ace* allele frequency. The colored ranges show the selection coefficients corresponding to the 0.95 confidence intervals of the slope distribution, obtained by bootstrapping cages within each treatment group. The overlap between the orange (treated cages) and dark blue (untreated cages) restricts the possible values of *s* and *h*. Within that overlap, lower values of *h* better explain both sets of data, as shown by higher values of the joint likelihood calculated from the bootstrap distributions (see **Methods**). All results of statistical tests can be found in **Extended Data Tables 4-6**.

Surprisingly, the malathion treatment did not alter the seasonal population dynamics in the malathion-treated cages (**Fig. 3a**). In all cages, population sizes increased through the summer months until reaching maximum size and then slowly declined in late Fall as abiotic conditions worsened. This suggests that population size in all cages is primarily limited by high levels of pre-adult mortality, potentially masking the additional mortality imposed by malathion.

Consistent with our predictions, the frequency of the resistant *Ace* alleles increased rapidly in the malathion-treated cages, followed by a frequency-dependent decline after insecticide removal. By tracking the resistant *Ace* alleles, we found that the R2 and R3 allele frequencies increased during malathion selection (R2: t = 13.93, p < 0.001; R3: t = 27.59, p < 0.001) (**Fig. 3b**), with the speed of change consistent with the selection coefficients (*hs*) on the order of 150-200% per generation. In contrast, the R1 allele showed no significant increase during treatment, consistent with its lack of detectable malathion resistance in lab assays (**Fig. 3b**, **Fig. 2d**). After insecticide removal, only the most common resistant allele (R3) declined (t = -5.18, p < 0.001), which reflects fitness costs that became evident at high frequencies (**Fig. 3b, Extended Data Table 5**).

To understand how these allele frequency changes influenced population-level resistance, we tracked malathion resistance throughout the experiment. The frequency increase of the resistant *Ace* alleles coincided with a rapid rise in malathion resistance, with LD_50_ increasing from ∼2.2 ppm at the start of treatment to ∼12.6 ppm (t = 24.26, p < 0.001) (**Fig. 3d, Extended Data Fig. 3)**. Resistance subsequently declined to ∼10.6 ppm after malathion removal (t = -3.96, p = 0.001), consistent with fitness costs in the absence of selection (**Fig. 3d, Extended Data Fig. 3, Extended Data Table 4.1)**. To assess the contribution of the *Ace* locus in the evolution of resistance, we applied a viability selection model parametrized by lab-measured genotype viabilities (see **Methods**). This model closely recapitulated both allele frequency trajectories and resistance evolution, strongly suggesting that *Ace* is the major-effect locus driving rapid evolution of resistance to malathion selection (**Extended Data Fig. 4**).

In the absence of insecticide selection in the untreated cages, the resistant *Ace* allele frequencies remained stable supporting the prediction that their costs are hidden in heterozygosity (**Fig. 3c**). None of the resistant *Ace* alleles exhibited significant frequency changes (**Fig. 3c, Extended Data Table 6**), suggesting low selection coefficients (*s*) or dominance parameters (*h*) in the absence of malathion. Consistently, LD_50_ values remained low and unchanged throughout the experiment (∼2.2 ppm, t = 1.07, p = 0.288) (**Fig. 3d, Extended Data Fig. 3, Extended Data Table 4.2**), indicating that overall resistance levels were unaffected in the untreated populations.

To further obtain bounds on evolutionary parameters for the resistant *Ace* alleles in organophosphate-free conditions, we fit the allele frequency data of the treated and untreated cages to a viability selection model where all resistant alleles share the same fitness cost (see **Methods**). The model that fit both datasets best had low cost (*s* ∼ 2.5 to 5%) and low dominance values (*h* < 0.38) (**Fig. 3f**). At higher values of *h*, there is no single selection coefficient that can explain the patterns of allele frequency change for both treated and untreated cages, and the likelihood of the data is higher the closer *h* is to zero. This analysis confirmed our prediction that the fitness cost of the resistant *Ace* alleles is recessive in semi-natural conditions.

### Fluctuating insecticide selection generates chromosome-scale genomic perturbations at sites linked to the resistant *Ace* alleles

We then examined the genomic consequences of rapid frequency fluctuations at the *Ace* locus on linked genetic regions. Our experimental design presented us with a rare opportunity to study these effects directly because the linkage patterns of genome-wide SNPs with the *Ace* alleles at the beginning of the experiment are known from the inbred line genotypes. To characterize forward and reverse sweeps driven by the R2 and R3 alleles during and after malathion treatment, respectively, we used a two-step approach (see **Methods**). First, we employed quasibinomial logistic regression to estimate genome-wide allele frequency changes during malathion treatment (July 26 to September 21, 2021) and post-treatment (September 21 to December 22, 2021). Next, we quantified changes in allele frequencies in SNPs tightly linked to the sweeping R2 and R3 alleles across the genome (r2 > 0.03 and frequency > 60% in resistant lines, *Ace*-linked) and those that were largely unlinked (r2 < 0.01, control).

During malathion treatment, we observed a chromosome-wide forward sweep in the malathion-treated cages driven by the R2 and R3 resistant alleles (**Fig. 4a**). The sweep extended across the chromosome arms 3R (∼4-17Mbp), including the *Ace* locus (9.6Mb), and 3L (∼5-19Mbp), with *Ace*-linked sites showing consistently elevated effect sizes above control sites. The sweep’s width is consistent with the extremely rapid rise of the *Ace* alleles over 2-4 generations and the maximum recombination rate of ∼5cM/Mbp in *D. melanogaster*, implying the effects conservatively extending over a region of ∼5-10 Mbp. Our analysis also indicated the presence of two additional loci potentially responding to malathion selection. On chromosome arm 3L (∼19.5Mb), control sites showed a strong directional perturbation while *Ace-*linked sites uniformly decreased in frequency, suggesting interference from another adaptive locus with the *Ace* sweep (**Fig. 4a**, asterisk). A similar pattern was observed on chromosome arm 2R, where both *Ace*-linked and control sites showed comparable frequency perturbations, indicating a second putative locus under selection (**Fig. 4a** asterisk). The study of these additional adaptive loci to malathion treatment is beyond the scope of this work. As expected, in the untreated cages where the frequencies of the R2 and R3 alleles remained stable (**Fig. 3d**), *Ace*-linked sites showed no elevated frequency changes compared to the control sites (**Fig. 4b**).

**Figure 4.**
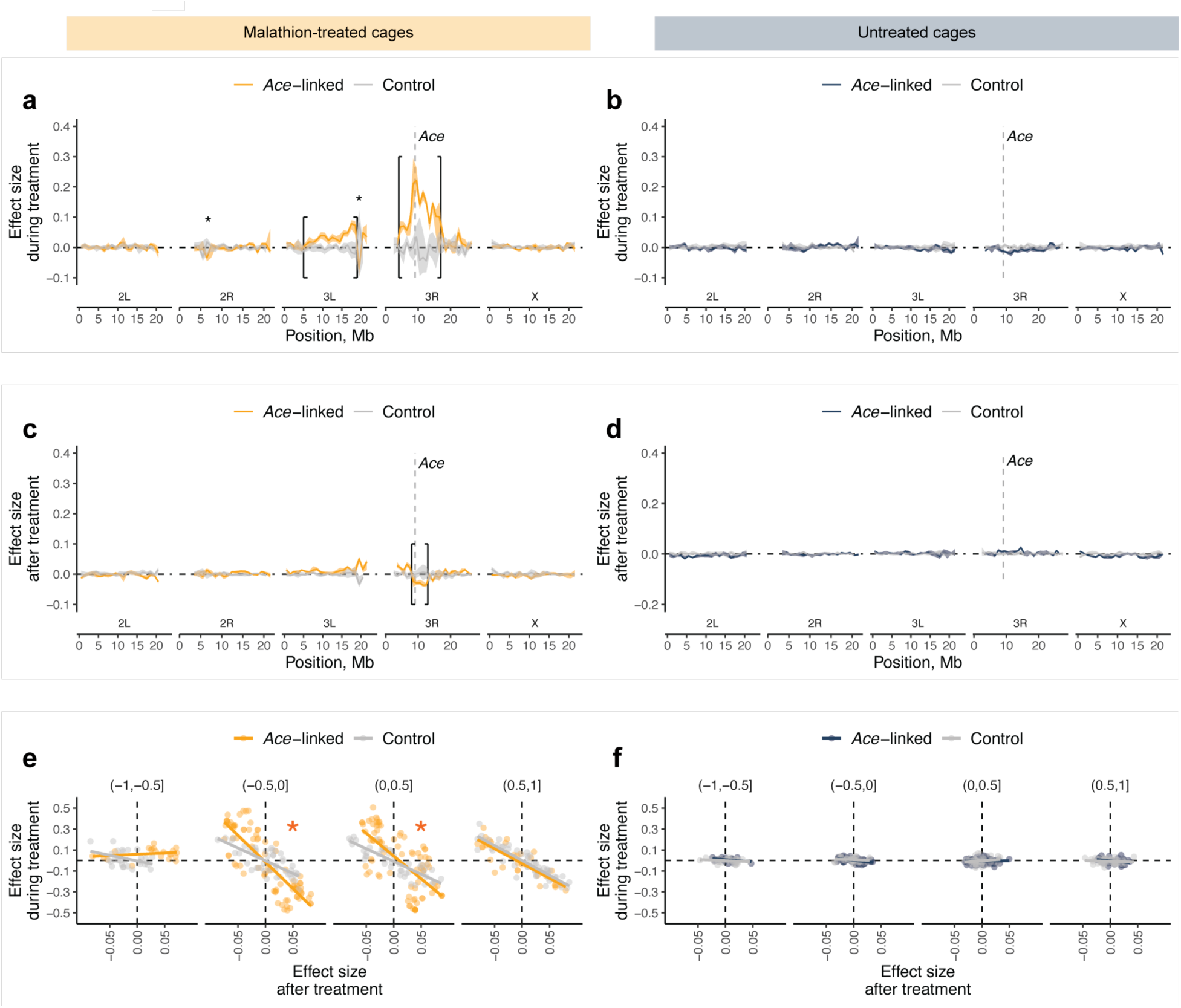
Strong fluctuating selection of resistant *Ace* alleles caused chromosome-scale forward sweep during malathion treatment and reverse sweep after treatment. **a-d**, Effect sizes (median allele frequency changes) for *Ace*-linked (orange/blue) and unlinked control (grey) SNPs in 1Mbp windows across the genome were estimated using quasibinomial logistic regression (see **Methods**). Changes during malathion treatment in treated (**a**) and untreated (**b**) populations. Changes after treatment cessation in treated (**c**) and untreated (**d**) populations. The brackets represent the inferred width of the sweep. Stars indicate additional adaptive loci to malathion treatment. **e-f**, Comparison of effect size for *Ace*-linked (orange/blue) and unlinked control (grey) SNPs during (y-axis) and after malathion treatment (x-axis) in malathion-treated (**e**) and untreated (**f**) populations within 1Mb from *Ace*. Each panel shows SNPs grouped by their distance from *Ace* (in Mb): [-1, -0.5], [-0.5, 0], [0, 0.5], and [0.5, 1]. In panel **e**, red stars indicate regions where the regression slopes significantly differ between *Ace*-linked and unlinked control SNPs.

After malathion treatment ceased and selection reversed, we detected an ∼5Mbp reverse sweep driven by the R2 and R3 alleles in the malathion-treated cages (**Fig. 4c**). Individual SNPs within 1Mbp of the *Ace* locus demonstrated a strong reversal effect, in which sites that increased in frequency during treatment tended to decrease afterwards, with this reversal being significantly stronger for *Ace*-linked compared to control sites (**Fig. 4e**). As expected, in the untreated populations, *Ace*-linked sites showed neither significant median frequency changes (**Fig. 4d**) nor the reversal behavior (**Fig. 4f**). Notably, recombination through the course of the experiment degraded the initial haplotype structure, suggesting the true extent of the reverse sweep may exceed 5 Mbp. Together, these findings confirmed our predictions that large-amplitude frequency fluctuations of adaptive alleles under temporally varying selection can drive significant effects on linked genomic variation.

## Discussion

Our study provides compelling empirical evidence for beneficial dominance reversal maintaining a large-effect resistance polymorphism at the *Ace* locus under temporally fluctuating insecticide selection in *D. melanogaster*. Previous studies have suggested the presence of beneficial reversal of dominance of resistant *Ace* alleles in *D. melanogaster* and other resistant insect populations based on the extension of Wright’s physiological theory^20^, enzymatic studies^22,24^, and analysis of fitness-associated phenotypes^19^. While these approaches can provide insights into the functional effects of resistance mutations, they do not necessarily predict organismal and fitness consequences^31^. For example, our results with the R1 allele illustrate this limitation - while enzymatic studies predicted it would confer similar levels of resistance as the R2 allele to malaoxon (the active breakdown product of malathion)^22^, we found a very small resistance benefit to malathion at the organismal and fitness levels. The novelty of our work with the *Ace* locus in *D. melanogaster* lies in combining laboratory assays, field mesocosm experiments with insecticide manipulation, and mathematical modeling to directly measure selection and dominance effects on the organismal level and on fitness in semi-natural conditions.

The beneficial dominance reversal we observed at the *Ace* locus suggests this mechanism may be widespread in maintaining functional genetic variation at large-effect loci under temporally fluctuating selection. Indeed, a growing body of literature indicates that beneficial reversal of dominance acts as a common mechanism for maintaining large-effect genetic variation across different forms of antagonistic selection between environments (e.g., fitness components, sexes, or niches/environments)^10,11^. For example, one allele of the large-effect *RXFP2* locus in Soay sheep has a dominant beneficial effect on male mating success but a recessive deleterious effect on male survival^32^. Similarly, the late maturation allele of the large-effect *VGLL3* locus in Atlantic salmon is partially dominant and likely beneficial in females, yet recessive and likely deleterious in males^33^. In the context of temporally fluctuating selection, large-effect loci coding for protein polymorphisms (e.g., enzymes, transcription factors) that are specialized for alternative contexts/substrates may commonly display beneficial reversal of dominance, as environmental shifts render these variants functionally impaired in their unfavored conditions. This pattern follows naturally from Wright’s physiological theory of dominance^12^, which predicts that the loss of protein function in alternative environments leads to the shifting identity of the recessive, deleterious allele^10,12^.

Finally, our study examined the genomic consequences of beneficial reversal of dominance at the *Ace* locus under temporally fluctuating selection. Our field mesocosm experiment revealed substantial genomic effects, with the rapid frequency changes at the *Ace* locus driving both a chromosome-wide forward sweep during malathion treatment and a reverse sweep after treatment removal. These findings provide empirical validation of theoretical predictions that beneficial dominance reversal of large-effect alleles can generate large-amplitude frequency fluctuations with significant effects on linked genetic variation^2,29^. Given that the *D. melanogaster* genome contains a fair number (tens to hundreds) of large-effect alleles displaying large frequency shifts due to fluctuating selection^5,6^ and the likelihood that this is the case for many other species^34,35^, fluctuating selection at large-effect loci could act as a major force shaping levels of genetic variation in natural populations^29^.

## Acknowledgements

This project was supported by grants from the National Institute of General Medical Sciences of the National Institutes of Health (Award number 1K99GM143455-01 to M.K, Grant 5R01GM100366-08 and 5R01GM137430-02 to P.S., Award Number 5R35GM118165-07 to D.A.P), the Sarah Hotchkis Ketterer Graduate Fellowship from Stanford University (to A.S.L.), the National Science Foundation (Grant PRFB 2109407 to M.C.B.), and the Stanford Biology Summer Undergraduate Research Program (to Z.K.M. and A.H.). The authors thank Marcus Feldman for helpful discussions and assistance with modeling *Ace* allele frequency evolution, Danae Papadopetraki for assistance with statistical analyses of the phenotypic and genomic data, Axel Wiberg for feedback on statistical analyses of the genomic data, Emily Behrman for feedback on the day degree model for generation estimation, Jessica Rhodes for helpful discussions and assistance with statistical analyses of the DEST data, and Anastasia Andreeva for assistance with phenotyping. The authors are very grateful to Jean Villa, Jessica Rhodes, Meike Wittmann, Noah Whiteman, and Marcus Feldman for feedback that improved the earlier versions of the manuscript.

## Author Contributions

M.K was responsible for conceptualization, investigation, formal analysis, writing original draft, writing-review and editing, visualization, and funding acquisition; A.L. was responsible for formal analysis, visualization, and writing-review and editing; M.B. was responsible for formal analysis and investigation; E.L. was responsible for formal analysis, visualization, and writing-review and editing; S.G. was responsible for formal analysis; A.H., CT, and Z.K.M. were responsible for investigation; P.S. and D.A.P. jointly supervised the research. P.S. was responsible for conceptualization, investigation, formal analysis, writing-review and editing, and funding acquisition. D.A.P. was responsible for conceptualization, investigation, formal analysis, writing-review and editing, and funding acquisition.

## Methods

### Population genomic analysis for variation of organophosphate-resistant *Ace* alleles across space and time

We first conducted a literature review to compile all known naturally segregating target site resistance mutations in the acetylcholinesterase (*Ace* gene) protein of *D. melanogaster* that confer resistance to organophosphates. Mutero and colleagues in 1994^36^ first identified the resistance mutations F115S, I199V, I199T, G303A and F368Y, and numbered them based on the precursor acetylcholinesterase (AChE) protein. Menozzi and colleagues in 2004^22^ identified combinations of the resistance mutations I161(199)V, G265(303)A, F330(368)Y and G368(406)A in worldwide organophosphate-resistant *D. melanogaster* strains and numbered them based on the mature AChE protein (numbers in the brackets refer to the precursor AChE protein). They did not identify the resistance mutation F77(115)S. At position 77, the mutation TTC to T**C**C changes phenylalanine to serine (Chr3R: 9,070,101, v5.39). At position 161, the mutation ATC to **G**TC changes the isoleucine to valine (Chr3R:9,069,721, v5.39). At position 265, the mutation GGC to G**C**C changes the glycine to alanine (Chr3R:9,069,408, v5.39). At position 330, the mutation TTT to T**A**T changes the phenylalanine to tyrosine (Chr3R:9,069,054, v5.39). At position 368, the mutation GGC to G**C**C/G**CT** changes the glycine to alanine (Chr3R: 9,063,921, v5.39). The AChE protein’s numbering changed between the studies after its crystallization in 2000, which defined the N-terminal signal peptide and the C-terminal sequence^37^. The mutation G368A was missanotated in the study of adaptation of *D. melanogaster* to organophosphate insecticides by Karasov and colleagues^23^.

### Organophosphate-resistant *Ace* alleles in DGRP and LNPA inbred lines

We characterized the presence of organophosphate-resistant *Ace* alleles in two fully sequenced inbred line panels, the Drosophila Genetic Reference Panel (DGRP) and our LNPA panel of inbred lines. The DGRP panel consists of 205 inbred lines derived from wild-caught females collected at the Raleigh, North Carolina Farmers Market in 2003^25^. The LNPA panel consists of 76 inbred lines derived from wild-caught individuals collected in Spring and Fall from Linvilla Orchards, Media PA USA^5^. Both panels contain the two common resistant *Ace* alleles (Ace-R1: <u>V</u>GFG and Ace-R3: <u>VAY</u>G) at high frequency, the sensitive *Ace* allele (Ace-S: IGFG), and other rare resistant alleles. The LNPA panel also contains the resistant *Ace* allele Ace-R2: <u>V</u>GF<u>A</u> at high frequency. We used the LNPA panel for our lab and field mesocosm experiments.

### Spatial and seasonal variation of organophosphate-resistant *Ace* alleles in natural *Drosophila melanogaster* populations

To examine the spatial variation of the I161V polymorphism, a marker of common organophosphate-resistant *Ace* alleles (Ace-R1: <u>V</u>GFG, Ace-R2: <u>V</u>GF<u>A</u>, Ace-R3: <u>VAY</u>G), we first analyzed the relationship between country-level pesticide use and I161V frequency in *Drosophila melanogaster* populations. We obtained pesticide use data from the Food and Agriculture Organization (FAO) FAOSTAT database^27^, which provides country-level estimates of pesticide use per area of cropland. For population genomic data, we used the Drosophila Evolution over Space and Time (DEST) database, which contains pooled genome-wide allele frequency estimates from 753 population samples of *D. melanogaster* collected worldwide. For our analyses, we included pesticide use data and genomic data for the period of 2000-2022.

To visualize the spatial distribution of insecticide use and frequency of the I161V polymorphism across locations, we created a choropleth map where countries are shaded based on their pesticide use per cropland area, with pie charts indicating the frequency of the resistant (I161V) and sensitive alleles at sampled locations. For each location, we calculated median allele frequencies across available samples for the period 2000-2022. To test whether insecticide use predicts the frequency of the resistance allele I161V, we fitted a linear mixed effects model using the lmer() function in the R package lme4 (version 1.1.33). The model included I161V frequency as the response variable and insecticide use per cropland area as a fixed effect. We included a random effect for cities.

In addition, to further test whether I161V shows evidence of spatial selection rather than drift, we compared its variance across samples collected in spring to matched control SNPs. We selected control SNPs based on three criteria: (1) located within 2Mb but at least 100kb away from *Ace* to avoid linked SNPs, (2) within the ln(3R)K cosmopolitan inversion to match inversion status, and (3) average frequency across samples within 4% of the median frequency of the I161V resistance mutation. We used the cumulative distribution function (CDF) to quantify how extreme the I161V variance was compared to the control SNPs.

To examine the temporal variation of the I161V polymorphism, we first tested two hypotheses: (1) whether the frequency of the resistant allele decreases during winter months when insecticide pressure is relaxed, potentially due to fitness costs, and (2) whether the frequency of the resistant allele increases during the growing seasons when insecticides are applied. We analyzed time series data from Linvilla Orchards, a non-organic orchard in Pennsylvania, USA. Samples were collected near apple trees from 2009 to 2015, with paired spring and fall collections each year, allowing us to test both hypotheses. To account for sampling variance in pooled sequencing data, we calculated the effective number of chromosomes sampled (Nc) following Machado et al. (2016), where Nc = (1/N + 1/R)^(-1), with N being the number of chromosomes in the pool (2 × number of flies) and R the read depth at the site. We then tested for systematic seasonal changes using generalized linear models (GLM) with binomial error structure. The models used the number of resistant and susceptible alleles (weighted by Nc) as the response variable and season as a categorical predictor. For the winter decline hypothesis, we paired each fall sample with the following spring sample. For the summer increase hypothesis, we paired each spring sample with the following fall sample from the same year. This approach allowed us to directly test for consistent directional changes in allele frequencies across seasons while accounting for both biological and technical sources of variance in the data.

Finally, to test whether the temporal dynamics of I161V reflect selection rather than drift, we compared its frequency fluctuations to those of matched control SNPs at Linvilla. To quantify temporal variation, we calculated the cumulative absolute change in I161V allele frequency over time as the sum of frequency differences between consecutive timepoints. This metric captures the total magnitude of frequency changes, regardless of direction. We selected control SNPs that had similar starting conditions as I161V, with initial frequencies matching the I161V frequency (0.42) at the first Linvilla timepoint within ±4%. To ensure a valid comparison, control SNPs were selected from within the same genomic region and inversion status as I161V, while excluding potentially linked sites (same criteria as for the spatial analysis). We then used the cumulative distribution function to assess whether I161V’s temporal variation was significantly greater than that observed in the matched control SNPs, which would suggest that its frequency changes are driven by seasonal selection rather than drift.

### Laboratory experiments to estimate the dominance of resistant *Ace* alleles for fitness-associated phenotypes in the presence and absence of malathion

To assess whether the dominance of each resistant *Ace* allele (Ace-R1: <u>V</u>GFG, Ace-R2: <u>V</u>GF<u>A</u>, Ace-R3: <u>VAY</u>G) over their sensitive counterpart (Ace-S: IGFG) reverses for fitness-associated phenotypes that are primarily under selection in organophosphate-rich and -free environments, we first used our LNPA panel of inbred lines to generate recombinant populations with homozygous sensitive, homozygous resistant, and heterozygous *Ace* genotypes for each *Ace* allele. This step was meant to bring potential deleterious, recessive mutations in the inbred lines into heterozygosity. We then measured malathion resistance, viability and fecundity in homozygous and heterozygous populations for each *Ace* genotype.

### Generation of recombinant populations with homozygous sensitive, homozygous resistant, and heterozygous *Ace* genotypes

To generate recombinant populations for each *Ace* allele, we first intercrossed 2-3 inbred lines with the same *Ace* allele using a round robin cross scheme (e.g., Line_1 SS x Line_2 SS, Line_2 SS x Line_3 SS, Line_3 SS x Line_1 SS) to bring all founders with the same *Ace* allele together (10 non-mated females x 10 males per round robin cross). We then combined 16 F1 flies (8 non-mated females and 8 males) from each of the round robin crosses per *Ace* allele to establish the next generation. Subsequently, we maintained the homozygous populations in vials and allowed them to recombine for four generations until we performed the phenotype measurements. We used lines 12LN6-70_(B33), 12LN6-92_(B3), LNPA-3 for the sensitive *Ace* allele (Ace-S: IGFG), lines 12LN6-86_(B29) and LNPA-53 for the R1 resistant *Ace* allele (Ace-R1: <u>V</u>GFG), lines LNPA-14, LNPA-17 and LNPA-87 for the R2 resistant *Ace* allele (Ace-R2: <u>V</u>GF<u>A</u>), and lines 12LN6-45_(B26), 12LN6-82_(B4), and 12LN6-94_(B19) for the R3 resistant *Ace* allele (Ace-R3: <u>VAY</u>G).

### Measurements of fitness-associated phenotypes

For all experiments, we first set up breeding cages with ∼ 500 4-to-7-day-old flies of homozygous or heterozygous populations for the sensitive and the resistant *Ace* alleles. To generate the heterozygous recombinant populations, we crossed SS non-mated females with homozygous males of each resistant allele (R1R1, R2R2, R3R3).

To compare the level of malathion resistance among the different alleles, we measured egg-to-adult viability on diets that ranged in malathion concentration from 0 ppm to 30 ppm. For this, we collected eggs of each population that were laid within a period of 24 hours and then introduced 50 eggs of each population into narrow vials containing approximately 7 ml of *Drosophila* growth media supplemented with the respective malathion concentration. We tested each of the malathion concentrations in duplicate vials. We used commercial malathion of 50% strength (Southern Ag Malathion 50% E.C.) and dissolved it in the *Drosophila* growth media. This medium was kept overnight at 4°C and used the next day for the assay. We monitored adult eclosion every evening over a period of 10 days until all adult flies had enclosed.

To compare viability among the different genotypes, we introduced 50 eggs of each population into narrow vials containing ∼7 ml of *Drosophila* growth media. Each of the genotypes were tested in 10 replicate vials. We monitored adult eclosion every evening over a period of 7 days until all adult flies had enclosed.

To compare fecundity among the different genotypes, we used density-controlled (∼50 eggs/vial) adult female progeny of each population. We introduced five 3-day-old females of each genotype in egg-laying bottles fitted with egg-laying caps filled with Drosophila growth media. Each day over a period of 3 days we counted the number of eggs laid. Each of the genotypes were tested in 8-10 replicate egg-laying bottles. The egg-laying caps were examined under a stereoscope where we counted the eggs laid each day over the 3-day period.

### Statistical analysis of fitness-associated phenotypes

We assessed fecundity, viability, and malathion resistance for the three genotypes (SS, SR, RR) of each *Ace* allele (R1, R2, R3) using R (version 4.2.2). For fecundity and viability, we fitted linear and logistic regression models, respectively, calculating estimated marginal means and performing pairwise comparisons adjusted for multiple testing using the Benjamini-Hochberg procedure. We then implemented a bootstrap procedure (1,000 resamples) to estimate deviation from additivity, selection coefficient, and dominance coefficient. The deviation from additivity was calculated as W[SR] - (W[SS] + W[RR]) / 2, where W represents fitness (fecundity or viability) for each genotype. The selection coefficient *s* was computed as (W[RR] – W[SS]) / W[SS], indicating the relative fitness of the resistant genotype compared to the sensitive genotype. The dominance coefficient *h* was calculated as (W[SR] – W[SS]) / (W[RR] – W[SS]), measuring the degree to which the heterozygote phenotype is closer to either homozygote. Significance was assessed by testing whether results differed from zero (additivity and selection) or 0.5 (dominance). For malathion resistance, we estimated LD_50_ values for each genotype using four-parameter logistic dose-response models f(x) = c + (d-c) / (1 + e^(b(x-e))), where e is LD_50,_ d (upper limit) = 100%, c (lower limit) = 0%, and b is the relative slope around e. For this, we used the L.4() function in the drc package (version 3.0.1)^38^ in R (version 4.2.2) with egg to adult viability at the range of malathion concentrations as the response variable. We then used a bootstrap procedure (1,000 resamples) to compare LD_50_ values between genotypes and calculate the same coefficients, substituting LD_50_ values for W in the formulas. For all analyses, bootstrap resampling was used to calculate confidence intervals and p-values, with significance assessed against null hypotheses of 0 (additivity and selection) and 0.5 (dominance).

### Field mesocosm experiments to track the evolution of malathion resistance and genome-wide allele frequencies in the presence and absence of fluctuating malathion selection

#### Experimental orchard and field mesocosm cages

To mimic the natural surroundings and weather conditions that flies are exposed to in nature, we conducted the study in an experimental orchard at the University of Pennsylvania in Philadelphia. The orchard consisted of cleared land with walk-in mesocosm cages (2m x 2m x 2m) constructed of fine mesh and built around metal frames (BioQuip PO 1406C). We planted a dwarf peach tree in each mesocosm cage to provide flies with shade. All peaches were removed before ripening to prevent flies from feeding or laying eggs on them. To provide flies with a feeding and breeding substrate, we added 1.5lb aluminum loaf pans with 400ml Drosophila growth media every two days during the experiment. To avoid an overcrowded developmental environment for the eggs, each loaf pan was covered with a screen mesh lid every two days during the experiment, and eggs were allowed to develop until eclosion ceased, at which point adult flies were released into the general population. The loaf pans were placed on a shelving unit oriented away from direct sunlight to protect them from rain and direct sun.

#### Establishment of experimental *D. melanogaster* populations in field mesocosm cages

To construct the baseline *D. melanogaster* population for the experiment, we crossed the 76 fully sequenced LNPA inbred lines from Pennsylvania. We used inbred lines to facilitate the use of haplotype inference to attain high effective sequencing coverage^39^. To initiate the baseline population, we combined 10 females and 10 males from each line into a single breeding cage. After 4 generations of mating and density-controlled rearing in laboratory conditions, we introduced 500 females and 500 males of a single age cohort into each mesocosm cage on June 21, 2021.

#### Malathion treatment over the course of the experiment

A diagram of the experimental design is shown in **Fig. 3a**. In the first step (*Establishment phase*: June 21 to July 20, 2021), we established 20 replicate populations in mesocosm cages and allowed them to expand in the absence of interventions. After establishment, the populations were split into ten malathion-treated and ten malathion-untreated populations. To mimic the insecticide exposure of flies in nature, the malathion-treated populations were exposed to malathion in the summer. In nature, insecticides are mainly applied during summer for pest control and their application ceases in the fall^40^. During the malathion treatment phase of the experiment, we sequentially exposed the populations to stepwise increasing concentrations of malathion, from 1.5 (July 21 to July 27, 2021) to 2.5ppm (July 28 to August 24, 2021) and 7.5ppm (August 25 to September 6, 2021). We used commercially available malathion grade of 50% strength (Southern Ag Malathion 50% E.C.) and dissolved it in the *Drosophila* growth media. We selected malathion doses based on the Malathion 50% E.C. label instructions to control small insect pests in the field and a previous long-term selection experiment for malathion resistance in *D. melanogaster*^41^. The medium supplemented with malathion was cooked once per week and stored at 4°C until deployed. After the malathion treatment, the experiment lasted until early winter (September 7 to December 22, 2021). Over the course of the experiment, we performed high-resolution temporal sampling (∼ bi-weekly sampling during the summer and monthly during the fall) to track evolution of malathion resistance and genome-wide allele frequencies.

### Tracking population size in the field mesocosms

To make relative comparisons of population sizes between the malathion-treated and untreated cages, we photographed 4 equal-size quadrats of the ceiling in each cage at dusk (approximately 2.5% of the ceiling for each quadrat). To count the number of adult flies in each photograph, we designed and validated an algorithm for semi-automated image processing. The number of adult flies in each of the 4 photographs was multiplied by 40 to correct for total surface area and obtain 4 population size estimates per cage.

#### Statistical analysis of population data

To assess the effects of time and malathion treatment on population size, we fitted a linear mixed-effect model with the estimated population size as the response variable. We defined our model to investigate the quadratic relationship of time with population size and to account for the random effects attributable to differences in cages. The regression analysis was performed in R (version 4.2.2) using lme4 (version 1.1.33).

### Tracking malathion resistance in the field mesocosms

To characterize the rate and direction of the evolution of malathion resistance in the malathion-treated and untreated populations, we repeatedly assayed the evolving populations. Prior to the malathion resistance assay, we collected F1 eggs from each mesocosm cage and transferred them in 2 groups of 200 eggs to bottles with media (i.e., density controlled) in a constant common garden environment (25°C, 12L:12D, 50% humidity) in the laboratory to distinguish evolution from phenotypic plasticity. We then performed the malathion resistance assay with density and age-controlled individuals from each mesocosm cage in the F2 generation (25°C, 12L:12D, 50% humidity). The common garden experiment for assaying malathion resistance took place at Stanford University with F1 eggs in bottles with media shipped overnight from the University of Pennsylvania.

#### Malathion resistance measurements

To characterize the level of malathion resistance in the malathion-treated and untreated populations, we tracked egg to adult survival in the F2 generation on diets that ranged in malathion concentration from 0 ppm to 15 ppm over the course of the experiment. For the collection of F2 eggs, we placed ∼ 400 4-to-7-day-old F1 individuals in breeding cages with egg lay caps for 5-7 hours. We then introduced 50 F2 eggs in a narrow vial containing ∼7ml of *Drosophila* growth media supplemented with malathion. We tested each of the malathion concentrations in duplicate vials. We used commercial malathion of 50% strength (Southern Ag Malathion 50% E.C.) and dissolved it in the *Drosophila* growth media. This medium was kept overnight at 4°C and used the next day for the assay. We monitored adult eclosion every evening over a period of 10 days until all adult flies had eclosed.

#### Statistical analysis of malathion resistance data

To assess the temporal dynamics of malathion resistance in both malathion-treated and untreated cages, we fitted separate linear mixed-effects models using the lmer() function in the R package lme4 (version 1.1.33). The response variable in these models was malathion resistance, measured as the LD_50_ value, the dose that causes 50% egg-to-adult mortality. For the malathion-treated cages, we modeled resistance changes over two distinct time periods: (1) from July 26, 2021, to September 21, 2021, when populations were evolving under malathion treatment, and (2) from September 21, 2021, to December 22, 2021, after malathion treatment had ceased. In contrast, for the malathion-untreated cages, we assessed resistance changes over the full experimental period, from July 26, 2021, to December 22, 2021. To avoid biases, we excluded data from July 13, 2021, because the LD_50_ estimate from this time point was underestimated due to a technical error. In each model, we included time point as fixed effect, while a random intercept was included for each cage to account for between-cage variability. The main outcome measure of malathion resistance for each cage at each time point was the mean of 1–2 technical replicate LD_50_ measurements. To estimate LD_50_ for each technical replicate per cage per time point, we fitted four-parameter logistic dose-response models using the L.4() function in the drc package (version 3.0.1) in R (version 4.2.2) with egg to adult viability at the range of malathion concentrations as the response variable.

### Track genome-wide allele frequencies in the field mesocosms over time

To track the trajectories of the resistant and sensitive *Ace* alleles and characterize the extent of the sweep caused by the *Ace* alleles, we tracked genome-wide allele frequencies of the evolving populations across the experiment using the software tool HAF-pipe. HAF-pipe estimates genome-wide allele frequencies in the evolving populations via inference of known founder haplotypes from low-coverage pooled sequencing data ^39^. As part of our experimental design, we performed whole-genome, low-coverage pooled sequencing of 100 randomly selected F1 females from each of the 20 evolving populations at eight time points through the experiment. To measure biological and technical variation, we performed whole-genome, low-coverage pooled sequencing of different sets of 100 randomly selected F1 females from the 20 evolving populations at two time points collected at Stanford (biological replicates) and of same samples of 100 randomly selected F1 females but processed independently (technical replicates). Last, to evaluate the accuracy of HAF-pipe, we performed whole-genome, high-coverage pooled sequencing of four biological replicate samples from the baseline population and for the evolving populations at an early and a late time point. The data for the untreated cages were also used for a recent independent study from Bitter and colleagues ^6^.

#### Sample collection, DNA extraction, library preparation, and sequencing

We collected samples from each population approximately every two generations during the summer phase and every generation during the fall phase. To collect the samples, we placed a new loaf pan with Drosophila growth media in each mesocosm cage one day before collection. The next day, we transferred five groups of 200 F1 eggs to bottles with media, where we controlled the density and maintained them in a constant laboratory environment of 25°C, 12L:12D, and 50% humidity. Three bottles with F1 eggs were kept at UPenn and two bottles with F1 eggs were shipped overnight to Stanford. At Stanford, F1 adults were allowed to lay eggs, and 200 F2 eggs were placed in vials for density control. For both the F1 and F2 generation, we harvested 4-to-7-day-old individuals after eclosion and preserved them in 100% ethanol in 1.5ml vials at -20°C until further processing.

For each sample, we prepared genomic DNA from pools of 100 ethanol-preserved females. We first submerged the samples twice in a rehydration buffer (400 mM NaCl, 20 mM Tris-HCl pH 8.0, 30 mM EDTA) for 15 min at room temperature and then isolated genomic DNA using the Monarch Genomic DNA purification kit (Cat No. 3010L, New England Biolabs). The manufacturer’s protocol was modified to include a step for tissue homogenization with two 3mm glass beads (Cat No. 11-312C, Fisher Scientific) in a bead beater for five minutes to ensure complete homogenization of the flies so that the genomic DNA from each fly was equally represented in the genomic DNA pool. We quantified the genomic DNA using the Qubit dsDNA BR (Broad-Range) Assay kit (Cat No. Q32853, Life Technologies) on a Qubit spectrophotometer and estimated its purity on a NanoDrop spectrophotometer.

We then prepared sequencing libraries using the Illumina DNA prep, (M) Tagmentation kit (Cat No. 20018705, Illumina Inc.) and the Illumina UD Indexes sets (Cat No 20027123, 20027214, 20042666, 20042667, Illumina Inc.) for unique dual indexing. We modified the manufacturer’s protocol by preparing libraries in 1/5 of the recommended reaction volume to maximize kit efficiency and pooling equimolarly amplified libraries in sets of eight after the library amplification step. After the cleanup, we quantified the pooled libraries using a Qubit spectrophotometer and an Agilent Bioanalyzer. Finally, the pooled libraries were 150bp paired-end sequenced on the NovaSeq 6000 S2 platform at the Sequencing Service Center at the Stanford Genomics Center and on a NovaSeq 6000 SP platform at Admera. All sequences will be deposited in SRA.

#### Processing of DNA sequencing data for genome-wide allele frequency estimation

We trimmed the raw sequencing reads of adapter sequences and bases with quality score < 20. We then mapped the trimmed reads to the *D. melanogaster* reference genome (v5.39) using bwa and default parameters ^42^, and aligned reads were deduplicated using Picard tools. We randomly sampled reads from the aligned bams to obtain an equivalent, genome-wide coverage of 8x across all samples. We computed haplotype-derived allele frequency (HAF) estimates using HAF pipe. Briefly, after using an expectation-maximization algorithm to estimate haplotype frequencies in overlapping genomic windows from mapped sequence reads (originally developed by Kessner and colleagues ^43^) at each site previously identified as polymorphic in the inbred reference panel, we summed the estimated frequencies of haplotypes carrying the alternate allele to obtain HAFs . We conducted haplotype inference in window sizes that varied proportionally to the expected length of un-recombined haplotype blocks, which varied as a function of the estimated number of generations since the original outbreeding. HAFs were compiled to only include those sites with an average minor allele frequency (MAF) > 0.02 in the baseline population, and present in at least one evolved sample at a MAF > 0.01.

### Track trajectories of sensitive and resistant *Ace* alleles in the evolving populations over time

We first estimated the frequency of each of the 76 founder haplotypes in the *Ace* region (Chr3R:9,063,921-9,069,721) in each of the malathion-treated and malathion-untreated populations at each time point, focusing on the founders containing the four known resistance mutations . To obtain an accurate measure of the frequency of each haplotype, we calculated the mean frequency across the 10 overlapping genomic windows that included each site within the region of interest.

We then estimated the relative frequency of each *Ace* allele, defined by the combination of the four known resistance mutations, in each population at each time point. Given our knowledge of the resistance mutation combinations for each founder, we computed the relative frequency of each *Ace* allele (Ace-S: IGFG, Ace-R1: <u>V</u>GFG, Ace-R2: <u>V</u>GF<u>A</u>, Ace-R3: <u>VAY</u>G, Ace-R4:IGF<u>A</u>, Ace-R5:IG<u>Y</u>G, Ace-R6:<u>VA</u>FG, Ace-R7: <u>VAYA</u>) by summing the frequencies of the haplotype blocks from the 76 founders that shared the same *Ace* allele. For founders heterozygous for the *Ace* alleles, we divided the contributions of their haplotype block evenly between the two corresponding *Ace* allele frequencies. *Ace* alleles with a frequency < 0.01% (Ace-R4:IGF<u>A</u>, Ace-R5:IG<u>Y</u>G, Ace-R6:<u>VA</u>FG, Ace-R7: <u>VAYA</u>) in the evolving populations were excluded from further analysis.

#### Statistical analysis of *Ace* allele frequency data

To assess the temporal dynamics of *Ace* allele frequencies (Ace-S: IGFG, Ace-R1: <u>V</u>GFG, Ace-R2: <u>V</u>GF<u>A</u>, Ace-R3: <u>VAY</u>G), we fitted separate linear mixed-effects models using the lmer() function in the R package lme4 (version 1.1.33). The response variables in these models were the frequencies of each *Ace* allele. For the malathion-treated cages, we modeled *Ace* allele frequency changes over two distinct time periods: (1) from July 26, 2021, to September 21, 2021, when populations were evolving under malathion treatment, and (2) from September 21, 2021, to December 22, 2021, after malathion treatment had ceased. In contrast, for the malathion-untreated cages, we assessed *Ace* allele frequency changes from the beginning of the treatment to the end of the experiment, from July 26, 2021 to December 22, 2021.

### Model *Ace* allele frequency evolution

We simulated the evolution of *Ace* allele frequencies and malathion resistance in our mesocosm experiments using a viability selection model^44^. Additionally, based on the changes in *Ace* allele frequencies in malathion-free environments across both sets of cages, we estimated the possible ranges for the fitness costs of and dominance of malathion resistance.

#### The impact of viability on *Ace* allele frequency dynamics and malathion resistance in the field mesocosm experiment

We investigated whether selection acting on viability recapitulates the allele frequency and malathion resistance dynamics observed in the field. We focused on viability as it is a key fitness-related trait in both malathion-rich and malathion-free environments. Furthermore, we had laboratory measurements of viabilities for *Ace* genotypes across ranges of malathion concentrations, allowing us to parametrize our model and test its predictions.

Within our model, the population frequencies of the *Ace* alleles change only due to selection that acts on viability. We assumed that the effects of other forces (mutation, recombination, and genetic drift) were negligible in comparison over the timescales that we considered (∼10 generations). Under these assumptions, the frequency dynamics of the sensitive (*f_0_*) and resistant (*f_1_*, *f_2_*, *f_3_*) *Ace* alleles follow from

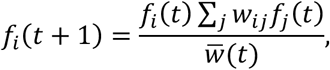

where *t* is time measured in generations, *W_ij_* is viability of individuals carrying both *i* and *j* alleles, and <u>*W*</u> is the mean population viability defined as

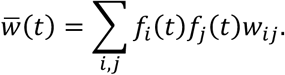

We used laboratory measurements to determine individual viabilities *W_ij_* in the absence and presence of malathion (at 2.5 and 7.5 ppm). We used the marginal mean values from the linear model for viability for each genotype and the predicted viability values from the dose-response analysis (see previous section on ‘Statistical analysis of fitness-associated phenotypes’). Due to the lack of measurements for heterozygous resistant genotypes, we assumed that combinations of resistant alleles are codominant, i.e. that *W_12_* = *0.5* · *W_11_* + *0.5* · *W_22_*; *W_13_* = *0.5 W_11_*+ *0.5* · *W_33_*; *W_23_* = *0.5* · *W_22_*+ *0.5* · *W_33_*. The initial *Ace* allele frequencies, *f_i_*(*0*), were estimated as the median values from the first sampling time point (July 13, 2021) for each treatment group.

To estimate *D. melanogaster* generation times at our eight sampling time points, we used a degree-day model based on Behrman and Schmidt^53^. This model assumes *D. melanogaster* requires 112 degree-days to complete one generation, with development occurring between 12°C and 29°C. We calculated daily degree-days as [(daily max. temp. + daily min. temp.) / 2] - 12°C, adjusting temperatures outside the 12-29°C range to these limits. Cumulative degree-days were tracked, with each 112 degree-day accumulation marking a new generation. This allowed us to estimate generations at each sampling time point throughout the experiment.

We then calculated the population resistance *R* as a weighted sum,

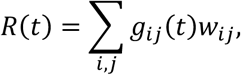

where *g_ij_*(*t*) are genotype frequencies and *W_ij_* are the corresponding viabilities. The initial genotype frequencies were estimated from the initial allele frequencies, assuming Hardy-Weinberg equilibrium:

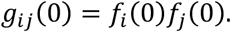

The consequent genotype frequencies followed from the viability selection model,

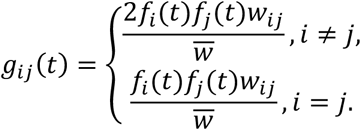

#### Inferring the ranges of possible fitness costs and dominance coefficients from empirical allele frequency trajectories

To estimate the fitness cost and dominance of malathion resistance in the malathion-free environment, we analyzed the empirical frequency trajectories of these alleles in both the untreated and treated populations after the removal of malathion. This approach allowed us to constrain the range of possible parameters that could explain the observed frequencies of the resistant alleles in both the treated and untreated populations. Assuming the resistant alleles share the same selection (s) and dominance (h) coefficients, and further that the cost of resistance is small (s<<1), the frequency dynamics follow

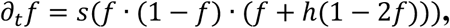

allowing us to find *s* that would be consistent with the observed slope *∂_t_f* of the frequency trajectory for a given *h*^54^.

To do that, we first bootstrapped the combined frequency trajectories of resistant alleles in individual cages within each treatment group 10,000 times. For each set of bootstrapped trajectories, we used linear regression (LinearRegression().fit function from the scikit-learn Python library^55^ to infer their slope. For a range of *h* ∈ [0,1], we then calculated the bounds on *s* by finding the values that correspond to the lower and upper bound of the 95% confidence interval of the bootstrap distribution. To further constrain the range of possible parameters, we similarly inferred the full distribution of *s* for each set of cages, and evaluated their overlap by computing the joint likelihood function,

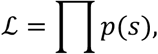

where *p*(*s*) is the probability density corresponding to a fixed value of *s*. Larger values of *L* therefore correspond to more likely combinations of *h* and *s* given the observed frequency dynamics.

### Characterize extent of forward and reverse sweep caused by *Ace* resistant alleles

To assess how fluctuating organophosphate selection on the R2 and R3 *Ace* resistant alleles affected genome-wide frequency trajectories of linked SNPs during and post-malathion treatment, we employed a two-step approach. First, we used quasibinomial logistic regression to estimate allele frequency shifts for genome-wide SNPs during malathion treatment (July 26 to September 21, 2022) and post-treatment (September 21 to December 22, 2021). Second, we developed a linked SNP selection procedure using linkage information from the inbred lines to identify SNPs linked (Ace-linked) and not linked (control) to the sweeping R2 and R3 *Ace* alleles across the genome. We used these sets of SNPs to quantify the extent of both the forward *Ace* sweep during malathion treatment and the reverse *Ace* sweep post-treatment.

#### Quasi-binomial logistic regression to estimate allele frequency differences in malathion-treated and untreated populations during and after the malathion treatment

To quantify allele frequency changes during malathion treatment (July 26 to September 21, 2022) and post-treatment (September 21 to December 22, 2021) in both treated and untreated populations, we used a quasibinomial logistic regression approach^5,48^. This method models the number of reads mapped to the alternative allele ‘α’ (relative to the reference allele ‘A’) at each biallelic SNP for each pooled sample. To account for both the expected accuracy of HAFs as well as pseudo-replication due to a fixed number of individuals in each sample ^49^, we estimated an effective read depth reads (at each biallelic SNP for each pooled sample per population using

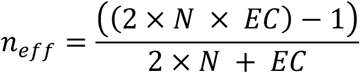

where *N* is the number of individuals per sample (100) and *EC* is the expected effective read coverage based on the accuracy of HAF-pipe.

We then used a logit link implemented in a quasi-binomial logistic regression to model the counts of the alternative allele ‘α’ (*C_α_* = *HAF* ∗ *neff*) and reference allele ‘A’ (*C_A_* = (*1* − *HAF*) ∗ *neff*) as a function of time in the malathion-treated and -untreated populations

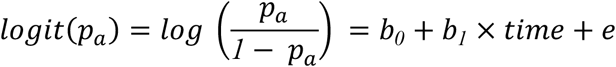

where *p_a_* is probability of observing the allele ‘α’, while the coefficients *b*_0_ and *b*_1_ correspond to the intercept and the effect of time, respectively. We fitted the model using the glm() function in R (version 4.2.2) with family = ‘quasibinomial’. To compute the allele frequency change (effect size) between two time points (*x*_1_ and *x*_2_), we calculated

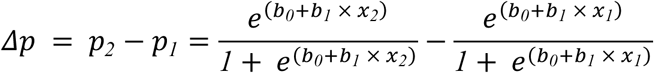

#### Analysis of allele frequency changes of Ace-linked SNPs

To quantify the extent of the forward and reverse *Ace* sweeps, we identified SNPs linked to the sweeping R2 and R3 *Ace* alleles (*Ace*-linked), as well as unlinked (control) SNPs, using whole genome data from the 76 sequenced Pennsylvania inbred lines (LNPA). First, we labeled R2 and R3 *Ace* alleles as “R” and the remaining alleles (R1 and S) as “S”, creating a binary “R”/“S” *Ace* locus across the inbred lines. Among the inbred lines, 54 were homozygous “S”, 5 heterozygous “R/S”, and 17 homozygous “R”. To select linked SNPs, we used the inbred lines data to compute r^2 between the binary “R”/“S” *Ace* locus and every reported biallelic SNP in the inbred LNPA lines as the squared correlation coefficient of allelic indicator variables^50^. We initially retained SNPs with r2 > 0.03, a threshold frequency previously used to deem two loci highly linked^6^. However, to avoid issues with low-frequency SNPs in finite populations, we added an allele count condition. We selected SNPs with r2 > 0.03 that also have an allele present in ≥60% of “R/R” lines and in ≤40% of “S/S” and “S/R” lines. This ensures selection of high-frequency, well-assorted SNPs between “R” and “S” lines, which are more informative for sweep analysis. This approach yielded 5672 Ace-linked SNPs across chromosomes 2, 3, and X. The 60% threshold was chosen as the highest value providing enough Ace-linked SNPs to uniformly cover the genome, with consistent results when varied. Additionally, for each Ace-linked SNP, we selected an unlinked control SNP with r2 < 0.01 with the binary “R”/“S” *Ace* locus. To ensure comparable initial conditions, unlinked control SNPs were chosen within 500kb of their linked counterpart and had an allele frequency within 0.05 of the Ace-linked SNP at the start of malathion treatment in each cage. Finally, to ensure that we track the SNP alleles that are linked specifically with the resistant lines, we transformed the alleles frequencies based on the sign of r — the non-squared correlation coefficient of allelic indicator variables. If r between *Ace* and a SNP is positive, then the alternative allele is assorted together with the resistant *Ace* alleles in the inbred lines, if the sign of r is negative, then the reference allele is linked to resistant *Ace* alleles, and in this case the alternative/reference labels were flipped, and the allele frequencies f_t were transformed to 1-f_t. This process yielded 4793 matched unlinked control SNPs, fewer than the linked SNPs due to the stringent matching criteria. To visualize the results, we aggregated SNPs into 1Mb windows along the genome and used bootstrap resampling to compute median GLM-estimated effect sizes and confidence intervals for Ace-linked and unlinked control SNPs separately for both forward and reverse sweeps. We excluded the areas around the centromeres and chromosome ends where less than 20 *Ace*-linked or control SNPs were observed. To characterize the width of each sweep, we performed a two-sample Mann-Whitney U-test within every 1Mb window across the chromosome and recorded the width of the contiguous range of windows around *Ace* in which the median for *Ace*-linked SNPs is significantly greater than the median for control SNPs at the 0.05 confidence level.

To quantify the reversal phenomenon during the *Ace* sweep post-malathion treatment, we analyzed allele frequency changes for the same Ace-linked and unlinked control SNPs during the post-treatment period (September 21 to December 22, 2021). We binned SNPs into 500Kb windows based on the genomic distance from the *Ace* locus on chromosome 3R and then compared post-treatment to during-treatment frequency changes by plotting them against each other within each bin. To quantify the reversal, we performed a joint linear regression for Ace-linked and control SNPs and recorded which bins have shown a slope for Ace-linked SNPs significantly more negative compared to control at the 0.05 confidence level.

### Data and Code availability

The data and associated code for this study will become available after manuscript acceptance in public repositories: Dryad (https://doi.org/10.5061/dryad.w0vt4b937) for the data and GitHub (https://github.com/mkarageorgi/Beneficial_Dominance_Reversal) for the code. The sequences of the LNPA inbred lines used to construct the baseline *D. melanogaster* population for the field mesocosm experiment are available on NCBI under accession ID PRJNA722305. The raw sequencing reads for pooled samples from the malathion-treated cages, collected during the field mesocosm experiment, will be deposited in the SRA BioProject under accession number PRJNA1031645. The raw sequencing reads for pooled samples from the untreated cages are already available in the same SRA BioProject (PRJNA1031645).

---

**Extended Data Figure 1.**
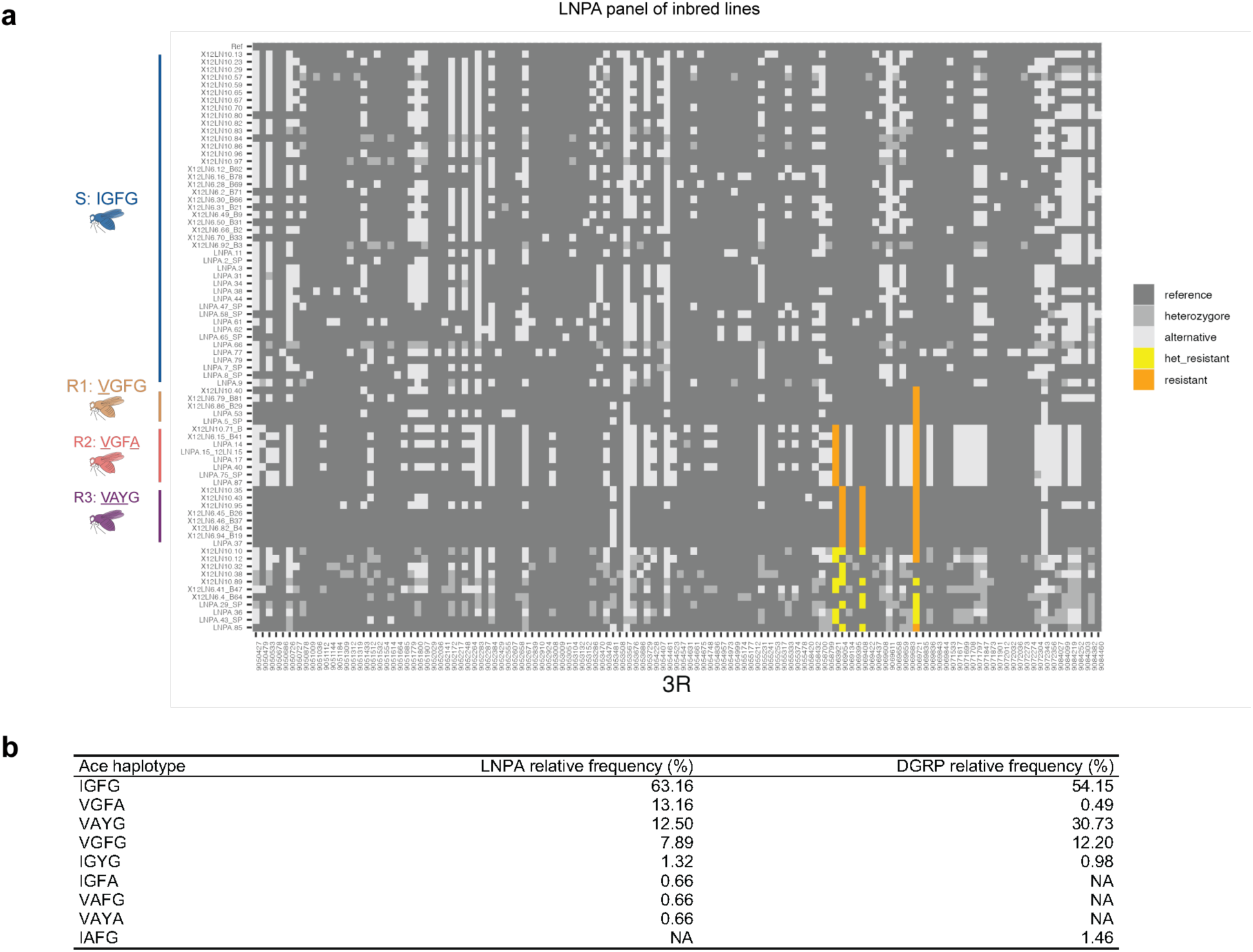
Sensitive and resistant *Ace* alleles in the panel of LNPA and DGRP inbred lines. **a,** Heatmap showing clustering of seventy-six *D. melanogaster* isogenic lines in the LNPA panel based on the sensitive or resistant *Ace* allele they carry (Ace-S: IGFG, Ace-R1: <u>V</u>GFG, Ace-R2: <u>V</u>GF<u>A</u>, Ace-R3: <u>VAY</u>G). The x-axis in the heatmap shows the genomic position on chromosome 3R around the *Ace* locus. Each row represents the *Ace* haplotype information of one isogenic line. Each column represents one biallelic exonic SNP at the *Ace* locus. Reference alleles are colored in dark grey, alternative alleles are colored in light grey, heterozygotes in medium grey. For the resistance mutations, alleles coding for the resistance mutations are colored in orange, alternative in grey, and heterozygotes in yellow. **b,** Table showing frequency of the sensitive and each of the resistant *Ace* alleles carrying different resistance haplotypes in the panels of LNPA and DGRP inbred lines.

**Extended Data Figure 2.**
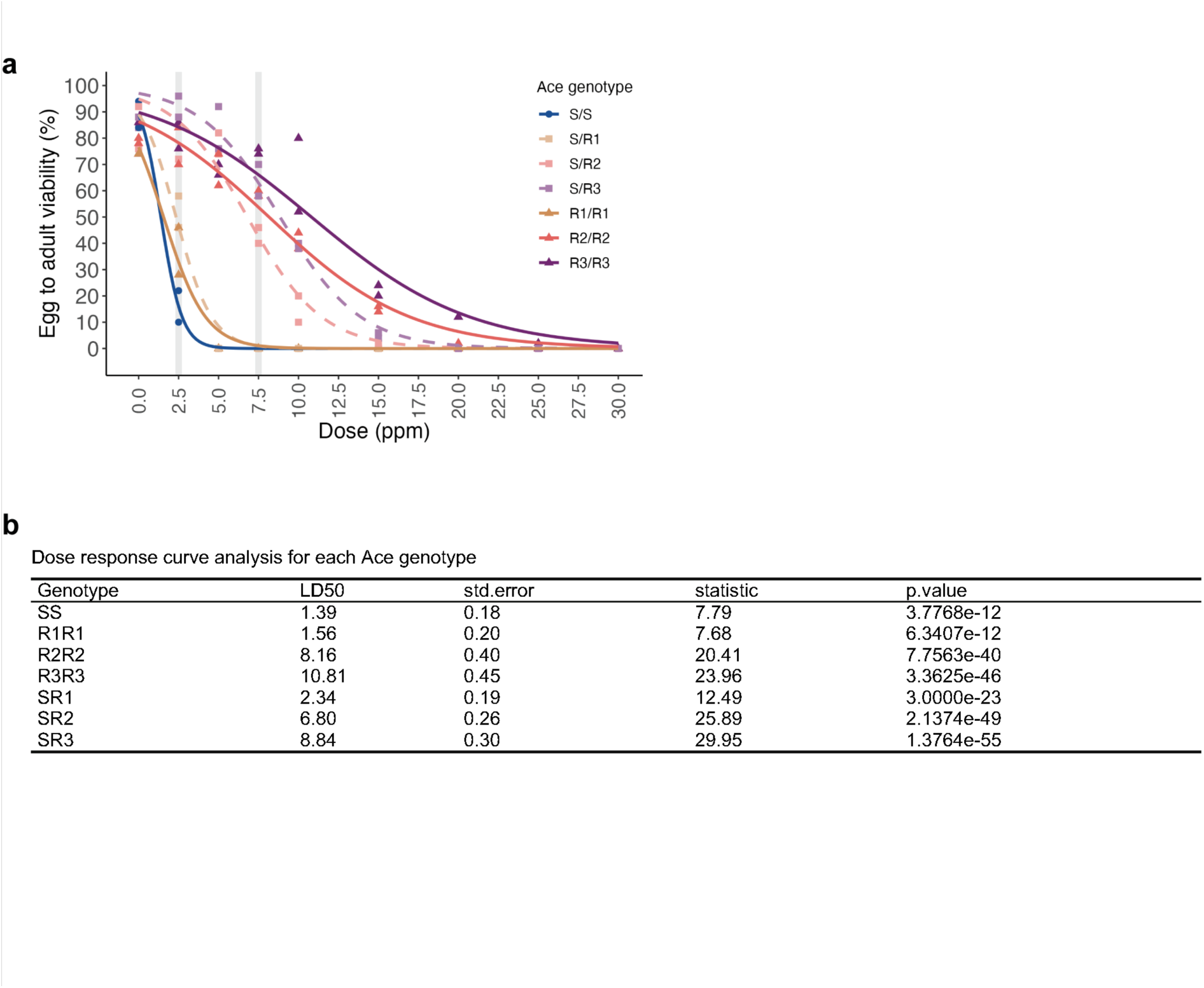
Comparison of malathion resistance among the *Ace* genotypes. **a,** Lab measurements of egg to adult viability in recombinant populations carrying different *Ace* genotypes on standard Drosophila media with a range of malathion concentrations (n= 2 technical replicates per genotype for each malathion doses, 50 eggs per replicate). Four-parameter logistic models fitted to data of each *Ace* genotype. The shaded areas highlight viability of each genotype at malathion doses (2.5ppm and 7.5 ppm) applied in the field mesocosm experiment. **b,** Table with estimated LD_50_ values for each *Ace* genotype from four-parameter logistic dose-response model (see also **Methods**).

**Extended Data Figure 3.**
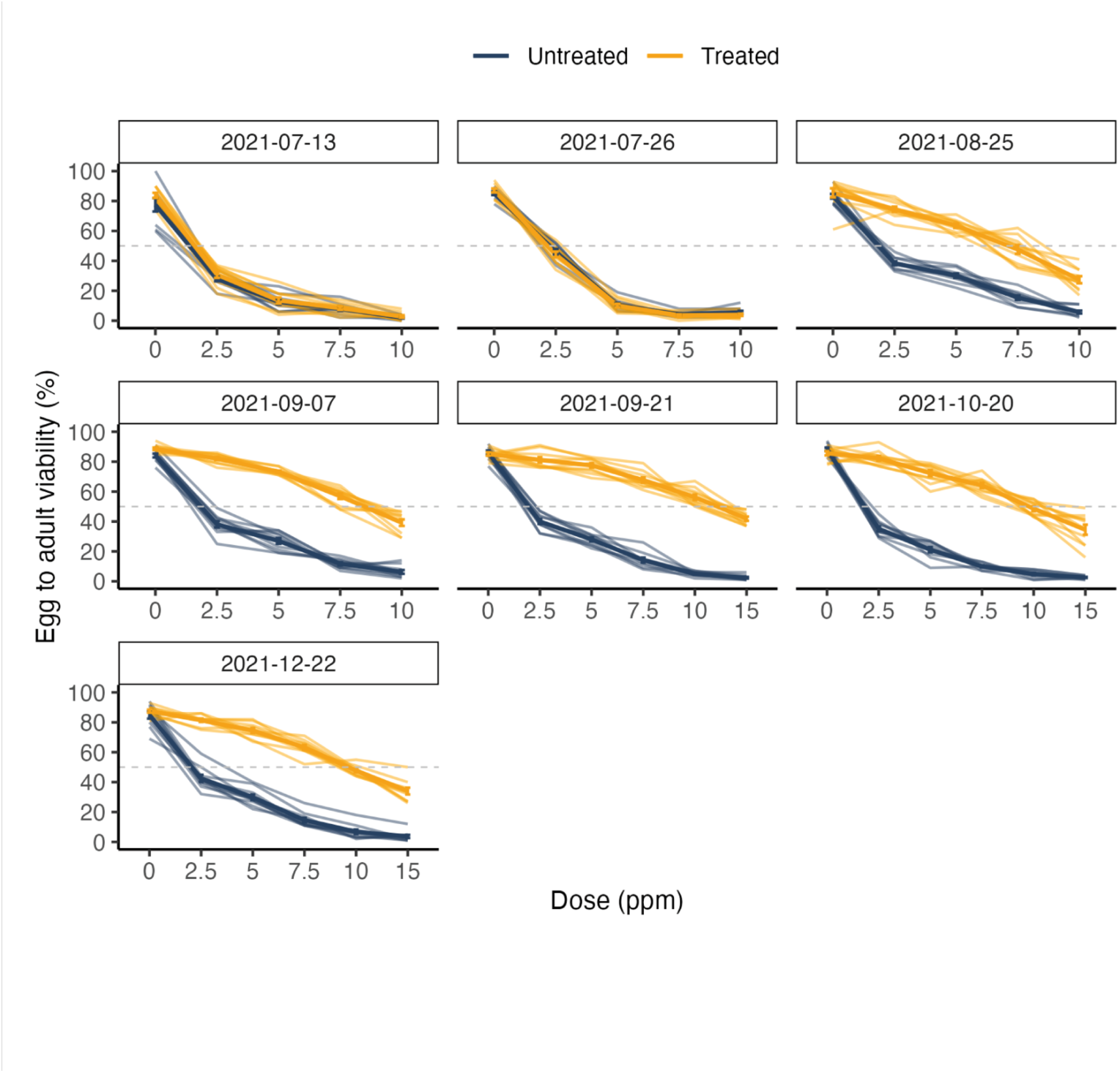
Tracking evolution of malathion resistance in the field mesocosm experiment. Egg-adult survival on diets with a range of malathion concentrations as measured after two generations of common garden rearing throughout the field mesocosm experiment. Light lines and points show the mean egg-adult survival for individual malathion-treated (orange) and untreated (blue) populations (n = 2 technical replicates, 50 eggs per replicate). Dark lines and points represent the mean ± s.e.m. from all populations for each treatment group (n = 10 populations per treatment group). Further information on experimental design can be found in **Methods**.

**Extended Data Figure 4.**
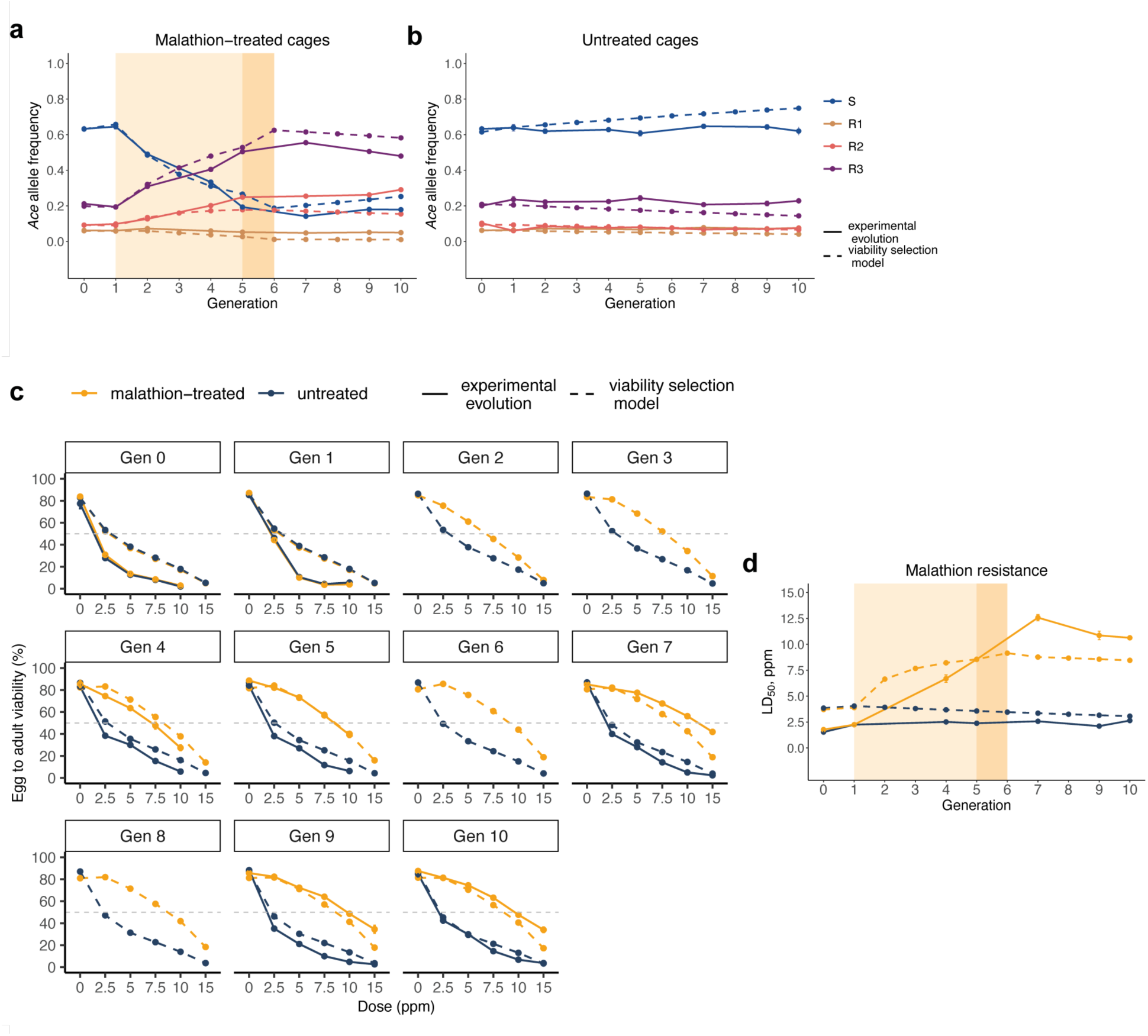
Modeling the impact of viability selection on *Ace* allele frequency dynamics and malathion resistance in the field mesocosm experiment. **a-b**, Comparison between the observed *Ace* allele frequency trajectories over 10 generations in the malathion-treated (**a**) and untreated (**b**) populations from the field mesocosm experiment (mean ± s.e.m.) and the expected trajectories from simulations based on a viability selection model using laboratory-derived parameters. See Methods for details about the viability selection model. **c-d**, Comparison between the observed egg-to-adult survival across a range of malathion concentrations (**c**) and the evolution of malathion resistance (LD_50_, ppm) (d) over 10 generations in the field mesocosm experiment (mean ± s.e.m.), and the expected results from the same model simulations. Dashed lines represent the expected values from the viability selection model, while solid lines represent the observed data from the field experiment. Generations were estimated based on a degree-day model starting from the first sampling point (July 13, 2021). In all plots, points correspond to each generation and estimated time point, and are connected by lines for visualization.

**Extendend Data Table 1.**

| Parameter | LD50 | Std.Error | P.value |
| --- | --- | --- | --- |
| SS mean | 1.39 | 0.19 | - |
| SR1 mean | 2.34 | 0.13 | - |
| R1R1 mean | 1.56 | 0.14 | - |
| SS vs. SR1 | -0.93 | 0.34 | 0.0063 |
| SS vs. R1R1 | -0.17 | 0.37 | 0.646 |
| SR1 vs. R1R1 | 0.76 | 0.39 | 0.0516 |
| SR1 - (SS + R1R1)/2 (additivity) | 0.76 | 0.32 | 1e-16 |
| (R1R1 - SS) / R1R1 (selection, s) | -0.25 | 0.29 | 0.751 |
| (SR1 - SS) / (R1R1 - SS) (dominance, h) | -1.29 | 5.10 | 0.751 |
| SR2 mean | 6.80 | 0.24 | - |
| R2R2 mean | 8.16 | 0.36 | - |
| SS vs. SR2 | -5.38 | 0.36 | 0 |
| SS vs. R2R2 | -6.72 | 0.42 | 0 |
| SR2 vs. R2R2 | -1.34 | 0.45 | 0.003 |
| SR2 - (SS + R2R2)/2 (additivity) | 1.87 | 0.35 | 1e-16 |
| (R2R2 - SS) / R2R2 (selection, s) | 3.83 | 1.05 | 1e-16 |
| (SR2 - SS) / (R2R2 - SS) (dominance, h) | 0.78 | 0.06 | 1e-16 |
| SR3 mean | 8.84 | 0.32 | - |
| R3R3 mean | 10.81 | 0.49 | - |
| SS vs. SR3 | -7.41 | 0.36 | 0 |
| SS vs. R3R3 | -9.37 | 0.79 | 0 |
| SR3 vs. R3R3 | -1.96 | 0.80 | 0.0145 |
| SR3 - (SS + R3R3)/2 (additivity) | 2.59 | 0.48 | 1e-16 |
| (R3R3 - SS) / R3R3 (selection, s) | 5.64 | 1.45 | 1e-16 |
| (SR3 - SS) / (R3R3 - SS) (dominance, h) | 0.77 | 0.07 | 1e-16 |

**Extendend Data Table 2.**

| Parameter | Viability | Std.Error | Lower.CI | Upper.CI | P.value |
| --- | --- | --- | --- | --- | --- |
| SS mean | 0.89 | 0.01 | - | - | - |
| SR1 mean | 0.86 | 0.02 | - | - | - |
| R1R1 mean | 0.76 | 0.02 | - | - | - |
| SS vs. SR1 | 1.42 | 0.27 | - | - | 0.0702 |
| SS vs. R1R1 | 2.61 | 0.47 | - | - | <.0001 |
| SR1 vs. R1R1 | 1.84 | 0.30 | - | - | 4e-04 |
| SR1 - (SS + R1R1)/2 (additivity) | 0.03 | 0.02 | -0.02 | 0.07 | 0.214 |
| (R1R1 - SS) / R1R1 (selection, s) | -0.15 | 0.03 | -0.2 | -0.09 | 0.001 |
| (SR1 - SS) / (R1R1 - SS) (dominance, h) | 0.29 | 0.20 | -0.15 | 0.64 | 0.214 |
| SR2 mean | 0.87 | 0.02 | - | - | - |
| R2R2 mean | 0.74 | 0.02 | - | - | - |
| SS vs. SR2 | 1.28 | 0.25 | - | - | 0.205 |
| SS vs. R2R2 | 2.96 | 0.53 | - | - | <.0001 |
| SR2 vs. R2R2 | 2.31 | 0.39 | - | - | <.0001 |
| SR2 - (SS + R2R2)/2 (additivity) | 0.05 | 0.02 | 0.01 | 0.1 | 0.027 |
| (R2R2 - SS) / R2R2 (selection, s) | -0.17 | 0.05 | -0.28 | -0.09 | 0.001 |
| (SR2 - SS) / (R2R2 - SS) (dominance, h) | 0.17 | 0.15 | -0.15 | 0.45 | 0.027 |
| SR3 mean | 0.86 | 0.02 | - | - | - |
| R3R3 mean | 0.78 | 0.02 | - | - | - |
| SS vs. SR3 | 1.42 | 0.27 | - | - | 0.0702 |
| SS vs. R3R3 | 2.43 | 0.44 | - | - | <.0001 |
| SR3 vs. R3R3 | 1.72 | 0.29 | - | - | 0.0018 |
| SR3 - (SS + R3R3)/2 (additivity) | 0.02 | 0.02 | -0.02 | 0.06 | 0.286 |
| (R3R3 - SS) / R3R3 (selection, s) | -0.13 | 0.02 | -0.18 | -0.08 | 0.001 |
| (SR3 - SS) / (R3R3 - SS) (dominance, h) | 0.32 | 0.19 | -0.15 | 0.67 | 0.286 |

**Extendend Data Table 3.**

| Parameter | Fecundity | Std.Error | Lower.CI | Upper.CI | P.value |
| --- | --- | --- | --- | --- | --- |
| SS mean | 200.22 | 11.99 | - | - | - |
| SR1 mean | 165.33 | 14.01 | - | - | - |
| R1R1 mean | 136.50 | 14.86 | - | - | - |
| SS vs. SR1 | 34.89 | 19.81 | - | - | 0.14 |
| SS vs. R1R1 | 63.72 | 20.42 | - | - | 0.01 |
| SR1 vs. R1R1 | 28.83 | 20.42 | - | - | 0.17 |
| SR1 - (SS + R1R1)/2 (additivity) | -3.03 | 17.26 | -36.62 | 31.59 | 0.879 |
| (R1R1 - SS) / R1R1 (selection, s) | -0.32 | 0.09 | -0.48 | -0.12 | 0.003 |
| (SR1 - SS) / (R1R1 - SS) (dominance, h) | 0.55 | 0.40 | -0.07 | 1.32 | 0.881 |
| SR2 mean | 187.60 | 13.64 | - | - | - |
| R2R2 mean | 167.78 | 14.37 | - | - | - |
| SS vs. SR2 | 12.62 | 19.81 | - | - | 0.53 |
| SS vs. R2R2 | 32.44 | 20.33 | - | - | 0.37 |
| SR2 vs. R2R2 | 19.82 | 19.81 | - | - | 0.49 |
| SR2 - (SS + R2R2)/2 (additivity) | 3.60 | 18.42 | -33.15 | 43.16 | 0.834 |
| (R2R2 - SS) / R2R2 (selection, s) | -0.16 | 0.09 | -0.33 | 0.03 | 0.093 |
| (SR2 - SS) / (R2R2 - SS) (dominance, h) | 0.39 | 4.09 | -2.3 | 3.23 | 0.842 |
| SR3 mean | 202.00 | 12.71 | - | - | - |
| R3R3 mean | 106.44 | 11.99 | - | - | - |
| SS vs. SR3 | -1.78 | 17.47 | - | - | 0.92 |
| SS vs. R3R3 | 93.78 | 16.95 | - | - | <.0001 |
| SR3 vs. R3R3 | 95.56 | 17.47 | - | - | <.0001 |
| SR3 - (SS + R3R3)/2 (additivity) | 48.67 | 12.84 | 22.41 | 73.56 | 1e-16 |
| (R3R3 - SS) / R3R3 (selection, s) | -0.47 | 0.07 | -0.59 | -0.31 | 0.001 |
| (SR3 - SS) / (R3R3 - SS) (dominance, h) | -0.02 | 0.19 | -0.48 | 0.29 | 0.001 |

**Extendend Data Table 4.**
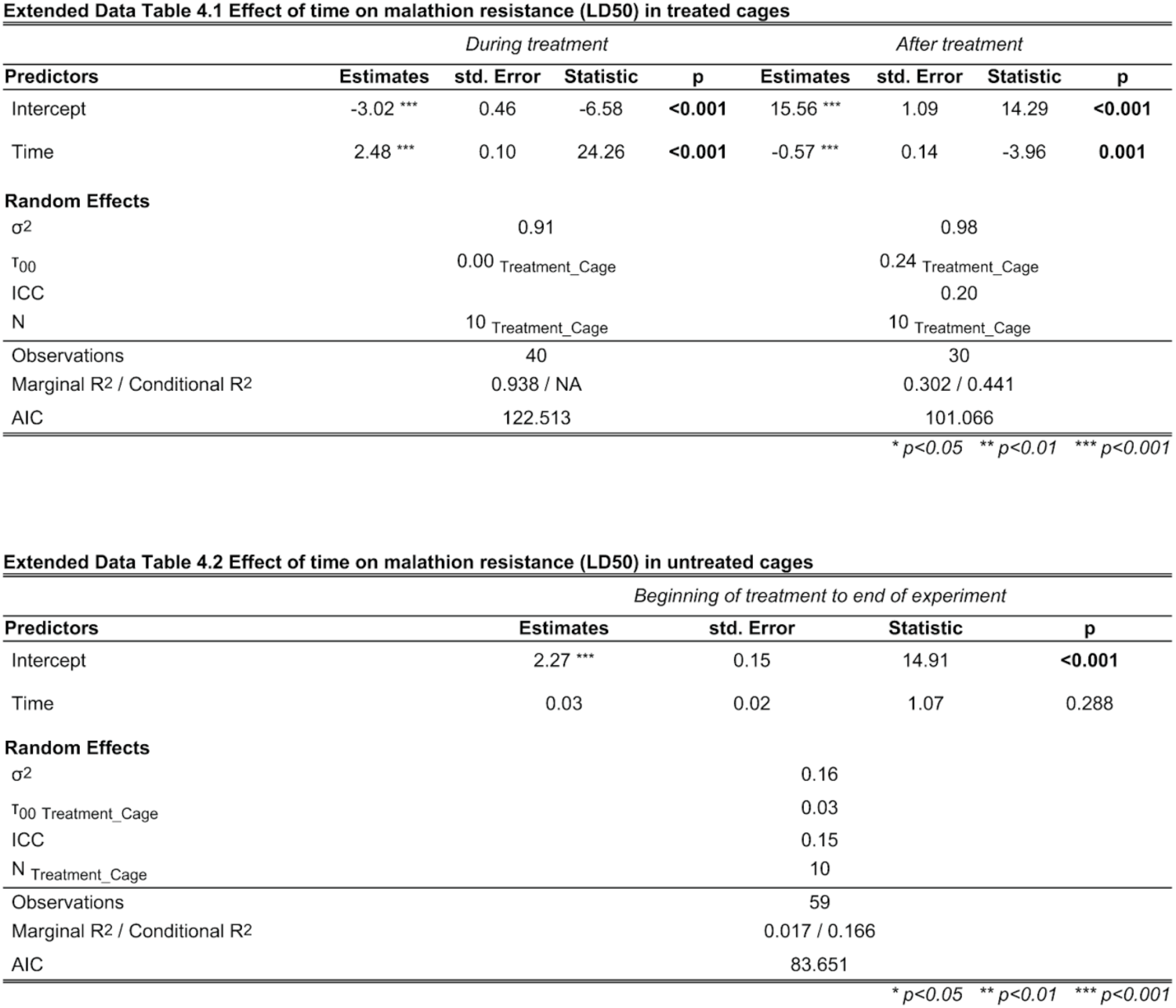

**Extendend Data Table 5.**
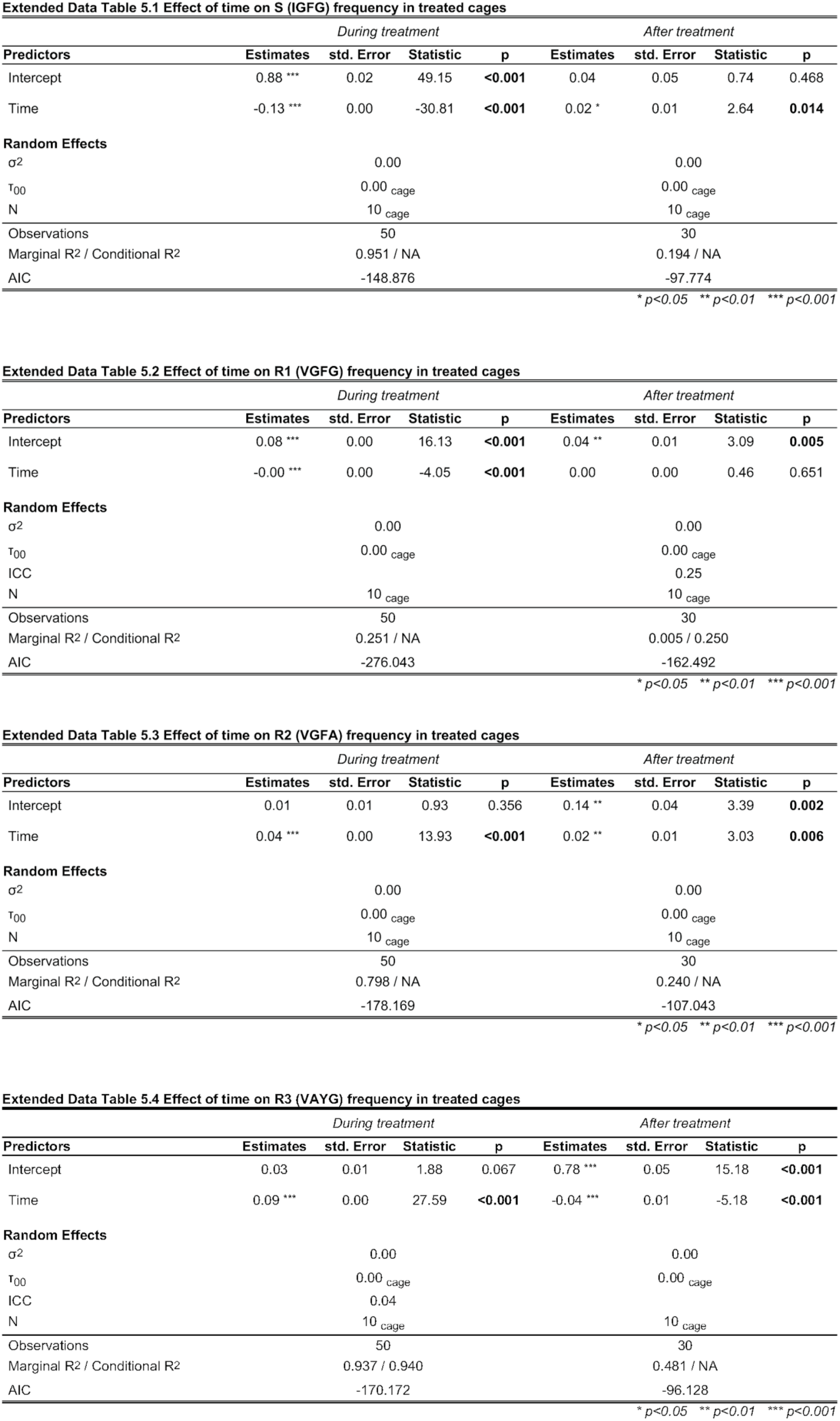

| Predictors | During treatment |  |  |  | After treatment |  |  |  |
| --- | --- | --- | --- | --- | --- | --- | --- | --- |
|  | Estimates | std. Error | Statistic | p | Estimates | std. Error | Statistic | p |
| Intercept | 0.88 *** | 0.02 | 49.15 | <0.001 | 0.04 | 0.05 | 0.74 | 0.468 |
| Time | -0.13 *** | 0.00 | -30.81 | <0.001 | 0.02 * | 0.01 | 2.64 | 0.014 |
| **Random Effects** |  |  |  |  |  |  |  |  |
| $\sigma^2$ | | 0.00 | | | | 0.00 | | |
| $\tau_{00}$ | | 0.00 cage | | | | 0.00 cage | | |
| N |  | 10 cage |  |  |  | 10 cage |  |  |
| Observations |  | 50 |  |  |  | 30 |  |  |
| Marginal R<sup>2</sup> / Conditional R<sup>2</sup> |  | 0.951 / NA |  |  |  | 0.194 / NA |  |  |
| AIC |  | -148.876 |  |  |  | -97.774 |  |  |

\* $p < 0.05$ \*\* $p < 0.01$ \*\*\* $p < 0.001$

| Predictors | During treatment |  |  |  | After treatment |  |  |  |
| --- | --- | --- | --- | --- | --- | --- | --- | --- |
|  | Estimates | std. Error | Statistic | p | Estimates | std. Error | Statistic | p |
| Intercept | 0.08 *** | 0.00 | 16.13 | <0.001 | 0.04 ** | 0.01 | 3.09 | 0.005 |
| Time | -0.00 *** | 0.00 | -4.05 | <0.001 | 0.00 | 0.00 | 0.46 | 0.651 |
| **Random Effects** |  |  |  |  |  |  |  |  |
| $\sigma^2$ | | 0.00 | | | | 0.00 | | |
| $\tau_{00}$ | | 0.00 cage | | | | 0.00 cage | | |
| ICC |  |  |  |  |  | 0.25 |  |  |
| N |  | 10 cage |  |  |  | 10 cage |  |  |
| Observations |  | 50 |  |  |  | 30 |  |  |
| Marginal R<sup>2</sup> / Conditional R<sup>2</sup> |  | 0.251 / NA |  |  |  | 0.005 / 0.250 |  |  |
| AIC |  | -276.043 |  |  |  | -162.492 |  |  |

\* $p < 0.05$ \*\* $p < 0.01$ \*\*\* $p < 0.001$

| Predictors | During treatment |  |  |  | After treatment |  |  |  |
| --- | --- | --- | --- | --- | --- | --- | --- | --- |
|  | Estimates | std. Error | Statistic | p | Estimates | std. Error | Statistic | p |
| Intercept | 0.01 | 0.01 | 0.93 | 0.356 | 0.14 ** | 0.04 | 3.39 | 0.002 |
| Time | 0.04 *** | 0.00 | 13.93 | <0.001 | 0.02 ** | 0.01 | 3.03 | 0.006 |
| **Random Effects** |  |  |  |  |  |  |  |  |
| $\sigma^2$ | | 0.00 | | | | 0.00 | | |
| $\tau_{00}$ | | 0.00 cage | | | | 0.00 cage | | |
| N |  | 10 cage |  |  |  | 10 cage |  |  |
| Observations |  | 50 |  |  |  | 30 |  |  |
| Marginal R<sup>2</sup> / Conditional R<sup>2</sup> |  | 0.798 / NA |  |  |  | 0.240 / NA |  |  |
| AIC |  | -178.169 |  |  |  | -107.043 |  |  |

\* $p < 0.05$ \*\* $p < 0.01$ \*\*\* $p < 0.001$

| Predictors | During treatment |  |  |  | After treatment |  |  |  |
| --- | --- | --- | --- | --- | --- | --- | --- | --- |
|  | Estimates | std. Error | Statistic | p | Estimates | std. Error | Statistic | p |
| Intercept | 0.03 | 0.01 | 1.88 | 0.067 | 0.78 *** | 0.05 | 15.18 | <0.001 |
| Time | 0.09 *** | 0.00 | 27.59 | <0.001 | -0.04 *** | 0.01 | -5.18 | <0.001 |
| **Random Effects** |  |  |  |  |  |  |  |  |
| $\sigma^2$ | | 0.00 | | | | 0.00 | | |
| $\tau_{00}$ | | 0.00 cage | | | | 0.00 cage | | |
| ICC |  | 0.04 |  |  |  |  |  |  |
| N |  | 10 cage |  |  |  | 10 cage |  |  |
| Observations |  | 50 |  |  |  | 30 |  |  |
| Marginal R<sup>2</sup> / Conditional R<sup>2</sup> |  | 0.937 / 0.940 |  |  |  | 0.481 / NA |  |  |
| AIC |  | -170.172 |  |  |  | -96.128 |  |  |

\* $p < 0.05$ \*\* $p < 0.01$ \*\*\* $p < 0.001$

**Extendend Data Table 6.**

| *Beginning of treatment to end of experiment* |  |  |  |  |
| --- | --- | --- | --- | --- |
| Predictors | Estimates | std. Error | Statistic | p |
| Intercept | 0.63 *** | 0.02 | 39.49 | <0.001 |
| Time | 0.00 | 0.00 | 0.21 | 0.838 |
| **Random Effects** |  |  |  |  |
| $\sigma^2$ | | 0.00 | | |
| $T_{00}$ cage | | 0.00 | | |
| ICC |  | 0.03 |  |  |
| $N_{cage}$ | | 10 | | |
| Observations |  | 69 |  |  |
| Marginal R<sup>2</sup> / Conditional R<sup>2</sup> |  | 0.001 / 0.032 |  |  |
| AIC |  | -196.762 |  |  |

\* $p < 0.05$ \*\* $p < 0.01$ \*\*\* $p < 0.001$

| *Beginning of treatment to end of experiment* |  |  |  |  |
| --- | --- | --- | --- | --- |
| Predictors | Estimates | std. Error | Statistic | p |
| Intercept | 0.06 *** | 0.00 | 14.68 | <0.001 |
| Time | 0.00 | 0.00 | 1.91 | 0.061 |
| **Random Effects** |  |  |  |  |
| $\sigma^2$ | | 0.00 | | |
| $T_{00}$ cage | | 0.00 | | |
| $N_{cage}$ | | 10 | | |
| Observations |  | 69 |  |  |
| Marginal R<sup>2</sup> / Conditional R<sup>2</sup> |  | 0.051 / NA |  |  |
| AIC |  | -370.061 |  |  |

\* $p < 0.05$ \*\* $p < 0.01$ \*\*\* $p < 0.001$

| *Beginning of treatment to end of experiment* |  |  |  |  |
| --- | --- | --- | --- | --- |
| Predictors | Estimates | std. Error | Statistic | p |
| Intercept | 0.07 *** | 0.01 | 9.08 | <0.001 |
| Time | -0.00 | 0.00 | -0.02 | 0.985 |
| **Random Effects** |  |  |  |  |
| $\sigma^2$ | | 0.00 | | |
| $T_{00}$ cage | | 0.00 | | |
| ICC |  | 0.02 |  |  |
| $N_{cage}$ | | 10 | | |
| Observations |  | 69 |  |  |
| Marginal R<sup>2</sup> / Conditional R<sup>2</sup> |  | 0.000 / 0.024 |  |  |
| AIC |  | -285.686 |  |  |

\* $p < 0.05$ \*\* $p < 0.01$ \*\*\* $p < 0.001$

| *Beginning of treatment to end of experiment* |  |  |  |  |
| --- | --- | --- | --- | --- |
| Predictors | Estimates | std. Error | Statistic | p |
| Intercept | 0.24 *** | 0.01 | 17.99 | <0.001 |
| Time | -0.00 | 0.00 | -0.86 | 0.392 |
| **Random Effects** |  |  |  |  |
| $\sigma^2$ | | 0.00 | | |
| $T_{00}$ cage | | 0.00 | | |
| ICC |  | 0.05 |  |  |
| $N_{cage}$ | | 10 | | |
| Observations |  | 69 |  |  |
| Marginal R<sup>2</sup> / Conditional R<sup>2</sup> |  | 0.010 / 0.064 |  |  |
| AIC |  | -222.940 |  |  |

\* $p < 0.05$ \*\* $p < 0.01$ \*\*\* $p < 0.001$

